# Insights into the Binding Mechanism of Ascorbic Acid and Violaxanthin with Violaxanthin De-Epoxidase (VDE) and Chlorophycean Violaxanthin De-Epoxidase (CVDE) Enzymes: Docking, Molecular Dynamics, and Free Energy Analysis

**DOI:** 10.1101/2020.06.07.138495

**Authors:** Satyaranjan Biswal, Parth Sarthi Sen Gupta, Haamid Rasool Bhat, Malay Kumar Rana

## Abstract

Photosynthetic organisms have evolved to work under low and high lights in photoprotection, acting as a scavenger of reactive oxygen species. The light dependent xanthophyll cycle involved in this process is performed by a key enzyme (present in the thylakoid lumen) Violaxanthin De-Epoxidase (VDE) in the presence of violaxanthin and ascorbic acid substrates. Phylogenetically, VDE is found to be connected with an ancestral enzyme Chlorophycean Violaxanthin De-Epoxidase (CVDE) present in the green algae on the stromal side of the thylakoid membrane. However, the structure and functions of CVDE were not known. In search of functional similarities involving this cycle, the structure, binding conformation, stability, and interaction mechanism of CVDE are explored with the two substrates in comparison to VDE. The structure of CVDE was determined by homology modeling and validated. In-silico docking (of first-principles-optimized substrates) revealed it has a larger catalytic domain than VDE. A thorough analysis of the binding affinity and stability of four enzyme-substrate complexes are performed by computing free energies and its decomposition, the root-mean-square deviation (RMSD) and fluctuation (RMSF), the radius of gyration, salt-bridge and hydrogen bonding interactions in molecular dynamics. Based on these, violaxanthin interacts with CVDE to the similar extent as that of VDE, hence its role is expected to be the same for both the enzymes. On the contrary, ascorbic acid has a weaker interaction with CVDE than VDE. As these interactions drive epoxidation or de-epoxidation process in the xanthophyll cycle, it immediately discerns that either ascorbic acid does not take part in de-epoxidation or this process requires a different cofactor because of the weaker interaction of ascorbic acid with CVDE in comparison to VDE.

## 1. Introduction

Without photosynethesis, which is triggered by light, plants and animals can barely exist on the planet. Due to varitions of light (both quantity and quality), photosynthetic organisms have developed different strategies to maintain a fine balance between light harvesting photochemistry and photoprotection[Külheim et al., 2002]. The photoprotection mechanism consists with the dissipation of excess light energy in the form of heat, thus avoiding photo-oxidative damages caused by the reactive oxygen species (ROS)[Siefermann-Harms, 1985; Stransky and Hager, 1970]. When the limit of light intensity exceeds, the photosynthetic light harvesting complex is regulated rather by non-photochemical quenching (NPQ) mechanisms responsible for dissipating excess absorbed light as heat[Foyer, 2018; Müller et al., 2001; Niyogi and Truong, 2013; Ruban et al., 2012]. The most exclusively and intensively investigated component of NPQ is the high energy state quenching (qE), which leads to dropping of pH in thylakoid lumen and acidification in excess light[Niyogi and Truong, 2013]. Among all, the dissipation of heat by carotenoids represents the most important photoprotective mechanism actively functioning in thylakoid membranes of plants and algae[Briantais et al., 1979; Britton, 1976; Bungard et al., 1999]. This process is accomplished by a cycle called “xanthophyll cycle” by interconverting violaxanthin carotenoid to antheraxanthin to zeaxanthin and vice-versa, respectively, at high light, low pH and low light, high pH conditions. Zeaxanthin is an antioxidant which scavenges reactive singlet oxygens by quenching of the excited singlet and triplet states of chlorophyll thereby accelerating the photoprotection capacity[Casper-Lindley and Björkman, 1998; Coesel et al., 2008; Lunch et al., 2013; Niyogi et al., 1997; Niyogi et al., 1998]. When light-driven proton translocation rate across the thylakoid membrane exceeds the dissipation rate of the proton gradient by ATPase leads to a reduction in pH in the thylakoid lumen activating Violaxanthin De-Epoxidase (VDE)[Baroli et al., 2003; Quaas et al., 2015; Yokthongwattana and Melis, 2006]. Activation of VDE occurs with a conformational change [Baroli et al., 2003; Bratt et al., 1995; Hager and Holocher, 1994; Morosinotto et al., 2002] and its association with the thylakoid membrane with the involvement of substrate violaxanthin [Bratt et al., 1995; Hager and Holocher, 1994; Morosinotto et al., 2002]. The role of zeaxanthin in photoprotection mechanism is vital as a mutant form of VDE shows an increased susceptibility to high light and reduced fitness in natural conditions due to de-excitation mechanisim[Külheim et al., 2002]. Whereas in low light conditions, the process gets reversed and zeaxanthin is converted back to violaxanthin by the stromal enzyme Zeaxanthin Epoxidase (ZE). [Bugos et al., 1998; Hieber et al., 2000]

A mutational variant (defective) of NPQ1 gene has been identified in the unicellular green alga Chlamydomonas reinhardtii and the Arabidopsis thaliana, which plays a significant role in the process of xanthophyll cycle and qE[Bugos et al., 1998; Hieber et al., 2000; Niyogi et al., 1998]. This NPQ1 mutant gene in Arabidopsis thaliana is unable to continue the process of VDE catalyzed conversion of violaxanthin (Vio) to zeaxanthin (Zea) in excessive intensity of light. On the other hand, in case of Chlamydomonas, the precise role of NPQ1 mutant in xanthophyll cycle is yet to be clearly understood. In case of Chlamydomonas genome, recent evidences suggest that it lacks a distinct orthologue of the VDE gene in plants and other algae [Anwaruzzaman et al., 2004; Hieber et al., 2000].

Recently, a new type of VDE gene has been found in Chlamydomonas reinhardtiien present at the stromal side of the thylakoid membrane, which helps translate a special type of VDE protein called Chlamydomonas CVDE (CrCVDE) protein. The evolutionary origins of plant-type VDE and CVDE are clearly distinct [Li et al., 2016]. For CVDE, CruP and CruA, there presents a common ancestor. CruA is known to be involved in bacterial carotenoid biosynthesis as a lycopene cyclase, whereas CruP is a paralogue of CruA, widely found in oxygenic photosynthetic living organisms[Bradbury et al., 2012]. Zhirong Li *et al*. hypothesized based on phylogenetic analysis that CVDE evolved by duplication of CruP from the ancestor of green algae and plants. They also categorized Chlamydomonas CVDE as a special type of protein which has been lost earlier. Billion years ago, it might be seen in some ancient clades of photosynthetic organisms such as green algae[Merchant et al., 2007], which effectively adapts itself to increase the photosynthetic efficiency and productivity.

Both VDE and ZE have been recognized as lipocalins, i.e. a member of multigenic protein family consisting of conserved structural pattern with a 8-strand b-barrel [Bugos et al., 1998; Hieber et al., 2000]. Lipocalin proteins are mostly involved in transporting small hydrophobic molecules such as steroids, bilins, retinoids, and lipids; one of the few lipocalins also exhibits catalytic activity in case of VDE and ZE[Hieber et al., 2000]. The structure of the VDE lipocalin domains (VDEcd and VDE central domain) possesses a structural pattern typical of that of a multigenic family, and also evidences a pH-dependent conformational change associated with protein activation that induces dimerization, allowing both Violaxanthin rings to react at the same time[Arnoux et al., 2009]. VDE has Cysteine-rich and Glutamate-rich two additional domains, as evident with no clear similarity with other proteins due to their peculiar amino acid composition[Bugos et al., 1998; Bugos and Yamamoto, 1996; Hieber et al., 2000; Morosinotto et al., 2002]. Both domains are found to be essential for protein activities, with their deletion leading to protein damage and inactivation of VDE [Bugos and Yamamoto, 1996]. While the Glutamine-rich domain has been suggested to be involved in protein association to the thylakoids membrane, the Cystine-rich one has unknown functional roles [Bugos and Yamamoto, 1996; Hieber et al., 2002]. While VDE is fairly known, studies on CVDE barely exist. There is no information about the sequence, structure, and functions reported for such enzyme giving any molecular level insights. As there is a phylogenetic connection between CVDE and VDE, it is instructive to predict their structural and functional similarities, exploring their interactions and binding mechanisms with substrates.

The aim of the present study is to investigate the molecular characteristics of the newly discovered de-epoxidase enzyme on the stromal side of the thylokoid membrane of Chlamydomonas CVDE (CrCVDE) protein and also inspect its roles in the xanthophyll cycle. A comparative assessment between VDE and CVDE is performed. The three dimensional (3-D) structural details of CVDE were deduced using a protocol that includes multiple-template homology modeling, protein threading *ab intio* modeling, and other Bioinformatics tools followed by structure validation. *In-silico* docking analysis of both enzymes CVDE and VDE were carried out with violaxanthin and ascorbic acid substrates to determine binding sites. Additional electronic structure information of the substrates were also provided. Post-molecular dynamics analysis, such as the root-mean-square deviation (RMSD) and fluctuation (RMSF), salt-bridges, h-bonds, key residues, principal components (PC) and free energies, demonstrates that both enzymes interact with violaxanthin to a similar extent, however, the interaction of ascorbic acid with CVDE is weaker than VDE. As the interaction of violaxanthin and ascorbic acid with VDE/CVDE guides the epoxidation or de-epoxidation process of the xanthophyll cycle, the latter substrate may not be required by CVDE or has trivial roles in the algae owing to the weak interaction.^31^

## 2. Materials and Methods

### 2.1. Sequence Analyses and Homology Modeling

The amino acid sequence of CVDE was retrieved from the protein sequence database UniPortKb with UniPortKb-ID: A0A2K3DUD0 (https://www.uniprot.org). The functional domains of both VDE and CVDE were identified by using Conserved Domains Database (CDD). BLASTp tool of NCBI was used for identifying a suitable template for the computational modeling of CVDE [Marchler-Bauer et al., 2010]. Multiple sequence alignment between template and target was performed by MultAlin tool (http://www.sacs.ucsf.edu/cgi-bin/multalin.py) and ESPript[Altschul et al., 1997]. The secondary structural elements were predicted using PSIPRED[Robert and Gouet, 2014]. The 3-D model structure of VDE enzyme (Chlamydomonas CVDE) was constructed using the multiple identity template module of homology modeling tool MODELLER v9.17[Buchan et al., 2013]. The best structure was selected according to the lowest discrete optimized protein energy (DOPE) score criteria and the side chain was further optimized using What if tool[Eswar et al., 2006].

The 3-D structure of VDE was retrieved from the protein data bank (PDB) with PDB ID - 3CQR. Both the refined CVDE and VDE structures were subjected to energy minimization using GROMACS [40]. The final energy-optimized model was validated by model validation server SAVeS (http://nihserver.mbi.ucla.edu/SAVES/). Further, for the model quality assessment and validation, PROCHECK [Vriend, 1990] and ERRAT [Abraham et al., 2015; Laskowski et al., 1993] were used. Energy profile characterization was performed using ProSA [Colovos and Yeates, 1993] and ProQ [Wiederstein and Sippl, 2007]. Analysis of bond lengths and bond angles was executed by MolProbity [Wallner and Elofsson, 2003].

### 2.2. pK_a_ Calculation

Multi-conformer continuum electrostatics (MCCE) v2.4 [Davis et al., 2007] and DELPHI v.4 [Song et al., 2009] were used to calculate pKa of residues in both VDE and CVDE at pH 7. Further, pK_a_ was calculated by PDB2PQR [Rocchia et al., 2002] server for comparison and a better accuracy. The PDB2PQR tool is integrated with several force field options, which was primarily designed for Adaptive Poisson-Boltzmann Solver (APBS) calculations. In the calculation process, structures were refined and reconstructed by fixing the missing atoms and several parameters through PDB2PQR like missing hydrogens, assigning atomic partial charges and radii from a specific force field. In the present pKa calculations, CHARMM force field was used [Dolinsky et al., 2007].

### 2.3. Quantum Calculation

Quantum simulation is needed to predict the correct geometry of small molecules like violaxanthin and ascorbic acid substrates in order to provide an accurate description of their binding mode and strength in the active sites of proteins. The Density functional theory (DFT) is presently the most successful approach to compute the electronic-structure properties [Patel et al., 2004]. Popular quantum mechanical descriptors such as the highest occupied molecular orbital (HOMO) and the lowest unoccupied molecular orbital (LUMO) play a major role in governing many chemical reactions[Arguelho et al., 2010; Singh et al., 2011]. Electronic descriptors that correlate to the physicochemical properties of molecules are energies of FMOs, band gap, total energy, dipole moment, ionization potential (IP), electron affinity (EA), electro negativity (χ), chemical potential (μ), global chemical hardness (η), global chemical softness (σ), electrophilicity index (ω), nucleophilicity (N), and hydrogen bond strength. Geometry optimization was carried out using Gaussian 16 at the level of DFT/B3LYP/6-31+G(d, p) followed by calculations of these DFT-based descriptors by post-processing of Gaussian output files of both the substrates (violaxanthin and ascorbic acid) with a home-built script[Eroğlu et al., 2007]. In assessment of charge densities on atoms in molecules and the optimized geometry, the hybrid DFT functional B3LYP is known to be fairly accurate[Kumari et al., 2014; Remko and von der Lieth, 2006]. Starting with the formatted check point files generated in geometry optimization, the charge density over meshes was calculated by CUBEGEN program to determine the molecular electrostatic potential (MEP) for both the substrates. GaussView [Dennington et al., 2007; Lee et al., 1988] program was used to visualize the molecular structure and partly to do analysis.

### 2.4. Molecular Docking

In view of comparing between the roles of VDE and CVDE in the xanthophyll cycle, understanding the details of binding of ascorbic acid and violaxanthin with CVDE and VDE is imperative. According to previous studies of the xanthophyll cycle, VDE is catalyzed by violaxanthin as well as ascorbic acid. This is conceptualized for CVDE in the current work to explore the binding mechanism. The structures of ascorbic acid and violaxanthin were obtained from Zinc database[Frisch et al., 2009] followed by geometry optimization by DFT. Using their optimized geometries, molecular docking was carried out with the protein enzymes by Auto Dock v4.2.3 program [Irwin and Shoichet, 2005]. A web based server 3-D Ligand site was used to determine grid values after identification of pocket binding residues [Morris et al., 2009]. Total 4 docking simulations were performed for two proteins with each of the two substrates. For accuracy, the torsional degrees of freedom, atomic partial charges, non-polar hydrogens and rotatable bonds of the substrates were taken into consideration while docking. A 65 × 65 × 65 grid box with spacing between points of 0.375 Å was used. Docking scores of substrate-protein complexes were analyzed by the Lamarckian Genetic Algorithm (LGA). The docked conformations were clustered with a 2 Å cut-off root mean square deviation (RMSD) and subsequently ranked according to their binding energy scores. PyMol was used for visualization and 2-D interaction plots were made by BIOVIA Discovery Studio Visualizer v4.5 (BIOVIA DSV) for the non-bonded interactions including hydrogen bond, van der Walls, π − π and other electrostatic interactions between the substrates and proteins. All of the four substrate-protein complexes were subjected to Molecular Dynamics thereafter for better understanding of interactions and stability of complex formation.

### 2.5. Molecular Dynamics (MD) Simulation of Complexes

GROMACS v2018 was used for carrying out molecular dynamics simulations to judge the stability, compact form, and molecular interactions of the complexes under dynamical situations at room temperature. The topology of the substrates was prepared using AnteChamber Python Parser interfaceE (ACPYPE) and PRODRG server [Wass et al., 2010]. The complexes were inserted in a cubical box containing extended simple point charge (SPC/E) water molecules and the GROMOS96 53a6 force field was used for the protein enzymes[da Silva and Vranken, 2012]. Counter ions (Na^+^/Cl^−^) corresponding to the physiological strength of 0.15 M were added to neutralize each simulation system. To avoid steric conflicts between atoms and high energy interactions, each electro-neutralized system was subjected to steepest descent energy minimization. After energy minimization, each system was treated with position restrained simulations in NVT and NPT ensemble. Each simulation consists of equilibration under NVT (canonical) ensemble for 1 ns at 300 K and subsequent 20 ns simulation under NPT ensemble during which the positions of the backbone of proteins were kept restrained. To constrain the covalent bonds, the Linear Constraint Solver (LINCS) algorithm was applied and electrostatic interactions were calculated using the particle mesh Ewald (PME) method. The cut-off radii for the coulomb and van der Waals interactions were fixed at 10.0 Å and 14.0 Å, respectively. The resultant MD trajectories saved at the interval of 100 ps were analyzed using the built-in modules of GROMACS and visual molecular dynamics (VMD 1.9.1)[Schuler et al., 2001]. 2-D plots depicting the intrinsic dynamical stabilities captured by the root mean square deviation (RMSD), root mean square fluctuation (RMSF), and radius of gyration (Rg) of the complexes were generated by Grace 5.1.23 program.

### 2.6. Principal Component Analysis (PCA)

Essential dynamics (ED) or principal component analysis (PCA) is an efficient statistical method that is applied to reduce the number of dimensions needed to describe protein molecular dynamics through a decomposition process [Humphrey et al., 1996]. PCA is a linear transformation that extracts the most important data using a covariance matrix or a correlation matrix (normalized PCA) modeled from atomic coordinates that describe the accessible degrees of freedom (DOF) of a protein, such as the cartesian coordinates that define atomic displacements or movements in each conformation in a trajectory. This covariance matrix is diagonalized to extract a set of eigenvectors and eigenvalues that reflect the correlated concert motion of the molecule [Amadei et al., 1993; Zhou et al., 2006]. The Gromacs module tool called g_covar was used to yield the eigenvalues and eigenvectors by calculating and diagonalizing the covariance matrix, whereas the g_anaeig tool was used to analyze and plot the eigenvectors[Swain et al., 2018].

### 2.7. Binding Free Energy Calculation

Molecular Mechanics/Poisson-Boltzmann Surface Area (MM/PBSA) is widely used for free energy calculation from MD trajectory. This method has been adopted to determine the binding affinity of substrate-protein complexes in many previous works in the post-molecular dynamics analysis[Van Aalten et al., 1995; Wang et al., 2018]. The binding free energy (ΔG_bind_) in a solvent medium was calculated as follows:

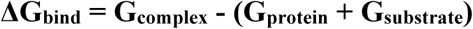

Where G_complex_ is the total free energy of the substrate-protein complex, G_protein_ and G_substrate_ are the total energies of protein and substrate alone in a solvent, respectively. The free energies for each individual G_complex_, G_protein_ and G_substrate_ were estimated by

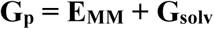

Where p can be protein, substrate, or complex. E_MM_ is the average molecular mechanics potential energy in vacuum and G_solv_ is the solvation free energy. The molecular mechanics potential energy was calculated in vacuum as follows:

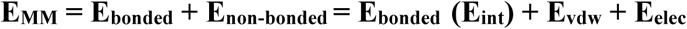

Where E_bonded_ or E_int_ is bonded interaction, which includes all bonded interactions like bond, angle, dihedral and improper interactions, and E_non-bonded_ is non-bonded interaction consisting of the sum of both van der Waals (E_vdw_) and electrostatic (E_elec_) interactions. E_bonded_ is always taken as zero. The solvation free energy (G_solv_) was estimated as the sum of electrostatic solvation free energy (G_polar_) and nonpolar solvation free energy (G_non-polar_)

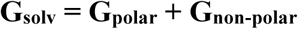

Where G_polar_, the polar solvation energy was determined and computed using the Poisson-Boltzmann (PB) linear equation and G_non-polar_ was estimated from the solvent-accessible surface area (SASA) as per the following equation:

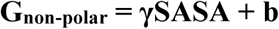

Where γ (coefficient related to surface tension of the solvent) = 0.02267 kJ/mol/Å^2^ or 0.0054 kcal/mol/Å^2^

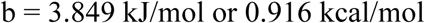

In this study, the binding free energies for all four complexes were calculated based on 1000 snapshots taken at an equal interval of time from 20 ns MD simulations. The per-residue energy contribution was also computed to understand the contribution of individual amino acids to the total binding energy.

## 3. Results and Discussion

### 3.1. Structural Overview and 3D Model Validation of CVDE

The conserved domain analysis shows that CVDE has a large domain of FAD binding-3 super family (PSSM ID-332373). This domain is involved in FAD binding in a number of enzymes. There are evidences that CVDE as hydroxybenzoate hydroxylase, a well-known member of this superfamily, exhibits catalytic behavior. Template identification through BLASTp search reveals a template, i.e. the crystal structure of 3-hydroxybenzoate hydroxylase (PDB ID-2DHK) with a sequence identity of 73% based on the target-template alignment (see Figure S1 of the Supporting Information). Employing Modeller v9.04, among the raw 3-D models developed, the model with the lowest DOPE score and RMSD value (on structural superposition on corresponding Cα atom pairs) with regard to the template structure is selected for further refinement. Loops were refined through Galaxy Loop refinement protocol, whereas the side chain improvement was performed using What-If. The refined model of CVDE was subjected to stereochemical quality assessment using various protein structure evaluation tools (see Table S1 and Figures S2, S3A-D) such as PROCHEK, Verify 3D, ERRAT, etc., and secondary sequence analysis by PSIPRED server (Figure S2). The accuracy of the model as per PROCHECK, integrated in Ramachandran plot, based on the dihedral (Phi and Psi) angles of amino acid residues is assessed to be 91.2% residues in the allowed region of the plot having an average G-factor score of 0.2 (Figure S3C). No residues are in the disallowed region of the Ramachandran plot which indicates quite high quality/accuracy of the model used (Figure S3C). Therefore, all the stereochemical properties of the model are within the acceptable range. The predicted model has a Verify3D score of 88.79 %, indicating the quality of the model is reasonably good as before. According to ProSA, the energy profile analysis (Table S1 and Figure S3D gives a Z-score of -4.54; ProQ analysis also indicates extremely good quality of the model based on their MaxSub (0.389 < 1.0) and LG scores (3.40 < 5.0). Bond length and bond angle analyses of the proposed models in MolProbity show none of the residues has bad side chains or main chain conflicts. Summarily, all the data for model validation are presented in Table S1 that contains as well the similar data for VDE (PDB ID-3CQR) for comparison. The overall analyses ensure the proposed model of CVDE is of good quality and can be considered for further studies. To assess the structural integrity of the modeled protein, architectural data on the 3-D structure analyzed with different tools are discussed next. Figures S3 represents the 3-D model structure and the topology of CVDE in solid-ribbon. The modeled CVDE structure has no identified barrel fold except thirteen α-helices together with twenty-three βstrands as seen in Figure S3A. The modeled structure consisting of 4 sheets (1 – parallel beta sheet, 1 - antiparallel-beta sheet, and 2 beta-alpha units with 61 residues in loop and 32 in helix); 8 beta hairpins, 12 beta bugles, 3-gama turns, 59 beta turns (29 h-bonds). Of 849 amino acids (aa) of the entire protein structure, 216 aa contribute to α-helices (24.65%), 292 aa to βstrands (33.23%), 5 aa (1.8%) and the rest 368 aa (42.09%), respectively, to turns and coils. In VDE (PDB ID-3qcr), the C-terminal pole of the α/βbarrel is found to be composed of loops with less than 5 aa over a short length. These are different from the loops at N-terminal pole which usually acts as an active binding site as evident in catalysis and other ligand binding mechanisms. Unlike VDE, CVDE does not have any specific evidence for its binding mechanisms through both the terminals, which urges for a deeper structural investigation to shed light on its catalytic properties.

### 3.2. DFT Studies: Electronic Descriptors

All the electronic descriptors which were obtained from DFT calculations of the substrates are presented in Table S2. Both the sign and magnitude of total energy express the likelihood of molecules[Dennington et al., 2007; Lee et al., 1988]. The total energy of ascorbic acid is -676.59 Hartree while violaxanthin has -1836.34 Hartree. The large negative values suggest that the molecules are energetically at their local minima having preferable configurations. The dipole moment of violaxanthin is 2.52 Debye while ascorbic acid shows the highest value of 4.04 Debye[Garbett and Chaires, 2012]. Elevated dipole moment enhances the hydrogen bond formation, nonbonding interactions, binding affinity and polar nature of a molecule[Lien et al., 1982]. The electron donating and receiving ability of a molecule can be determined by considering HOMO and LUMO energies of that molecule. Higher the energy of HOMO, higher will be its electron donating ability. It has been found that the energies of HOMOs are comparable for both compounds. Similarly, the LUMO orbital energies are also comparable. The LUMO energy level is affected by the presence of electron donating or withdrawing or both groups in the structure and is responsible for biological activity in addition to other factors. Thus, the study of frontier molecular orbitals (HOMO and LUMO) is quite useful. Chemical hardness (η) and softness (S) of a molecule can be determined from the LUMO — HOMO gap [Ayers et al., 2006]. A large gap is related to high kinetic stability and low chemical reactivity; a small gap is indicative of low chemical stability, because the addition of electrons to a high-lying LUMO and/ or removal of electrons from a low-lying HOMO is energetically favorable in any potential chemical reaction [Aihara, 1999]. In this study, ascorbic acid has the HOMO-LUMO gap of 0.35 eV while violaxanthin shows slightly a lower energy gap (0.29 eV). The larger chemical potential (−0.10 eV) along with higher chemical softness (6.84 eV) values may contribute to the higher chemical reactivity of violaxanthin than ascorbic acid (Table S2). The higher electronegativity (0.17 eV) and electrophilicity (0.09 eV) of ascorbic acid indicate its high susceptibility to withdraw electrons than violaxanthin.

### 3.3. Docking Analysis

It is well known that VDE binds with ascorbic acid and violaxanthin during the xanthophyll cycle. There are evidences that violaxanthin travels freely in the lipid phase of cell and does not bind with any pigment carrier proteins [2.18]. Therefore, the hunch is to find out whether these substrates can similarly work with CVDE. Figure 1 provides information about interactions present in the bonded complexes. In docking, the best docking pose is selected based on the highest docking score from the cluster analysis. Information about the highest docking scores, binding efficiency, and the number of h-bonds along with other interactional data for binding in each complex are given in Table 1. The VDE-ascorbic acid complex has binding affinity -7.4 kcal/mol, with four conventional h-bonds and two carbon-hydrogen bonds. In case of VDE-violaxanthin, there exist two conventional h-bonds. Despite a large number of h-bonds, VDE-ascorbic acid has a lower bindng affinity than VDE-violaxanthin (−9.14 kcal/mol). The two complexes of CVDE formed with ascorbic acid and violaxanthin have binding affinities -6.4 kcal/mol and -9.94 kcal/mol with the number of conventional h-bonds being three and five, respectively. Owing to the highest number of residues participating in the van dar Waals interaction, CVDE-violaxanthin has the strongest binding affinity among all. Despite slight variations, post-MD docking analyses also reveal the same trend in binding affinity among the substrate-enzyme complexes.

**Table 1:** Details about docking scores (binding energy) and interacting key residues of ascorbic acid and violaxanthin with the target enzymes VDE and CVDE.

| Enzymes | Ligands | Binding energy in kcal/mol (before MD) | Interactin g key residues | Types of bonding | Distance (Å) | Binding energy in kcal/mol (after MD) |
| --- | --- | --- | --- | --- | --- | --- |
| CVDE | Ascorbic acid | -6.40 | Ser137 | Conventional hydrogen | 4.85 | -5.90 |
|  |  |  | Ala 397 | Conventional hydrogen | 4.54 |  |
|  |  |  | Asp146 | Conventional hydrogen | 4.87 |  |
|  |  |  | Ser140 | Conventional hydrogen | 3.24 |  |
|  |  |  | Thr138 | Conventional hydrogen | 3.42 |  |
|  | Violaxanthin | -9.94 | Ser462 | Conventional hydrogen | 4.37 | -9.60 |
|  |  |  | Ala 529 | Pi-alkyl/alkyl | 4.24 |  |
|  |  |  | Leu 491 | Pi-alkyl/alkyl | 4.34 |  |
|  |  |  | Ala 463 | Pi-alkyl/alkyl | 4.55 |  |
|  |  |  | Ala 593 | Pi-alkyl/alkyl | 4.23 |  |
|  |  |  | Glu 493 | Conventional hydrogen | 4.41 |  |
| VDE | Ascorbic acid | -7.40 | Aln119 | Carbon hydrogen | 5.18 | -7.10 |
|  |  |  | Pro111 | Pi-alkyl/alkyl | 4.44 |  |
|  |  |  | Tyr198 | Conventional hydrogen | 5.85 |  |
|  |  |  | Asn110 | Conventional hydrogen | 4.00 |  |
|  |  |  | Thr112 | Conventional hydrogen | 2.48 |  |
|  | Violaxanthin | -9.14 | Thr112 | Conventional hydrogen | 2.29 | -9.40 |
|  |  |  | Ala115 | Pi-alkyl/alkyl | 4.49 |  |
|  |  |  | Ile105 | Pi-alkyl/alkyl | 5.48 |  |
|  |  |  | Asn110 | Conventional hydrogen | 4.50 |  |
|  |  |  | Ala211 | Conventional hydrogen | 3.91 |  |
|  |  |  | Val212 | Pi-alkyl/alkyl | 5.73 |  |
|  |  |  | Tyr198 | Pi-alkyl/alkyl | 6.48 |  |
|  |  |  | Cys118 | Pi-alkyl/alkyl | 4.48 |  |
|  |  |  | Ile105 | Pi-alkyl/alkyl | 4.30 |  |
|  |  |  | Ile243 | Pi-alkyl/alkyl | 5.52 |  |
|  |  |  | Val212 | Pi-alkyl/alkyl | 4.68 |  |
|  |  |  | Ala115 | Pi-alkyl/alkyl | 4.11 |  |
|  |  |  | Pro111 | Pi-alkyl/alkyl | 4.31 |  |
|  |  |  | Ala211 | Conventional hydrogen | 3.57 |  |
|  |  |  | Leu109 | Unfavorable donor-donor | 4.40 |  |
|  |  |  | Asn110 | Conventional hydrogen | 4.28 |  |

**Figure 1:**
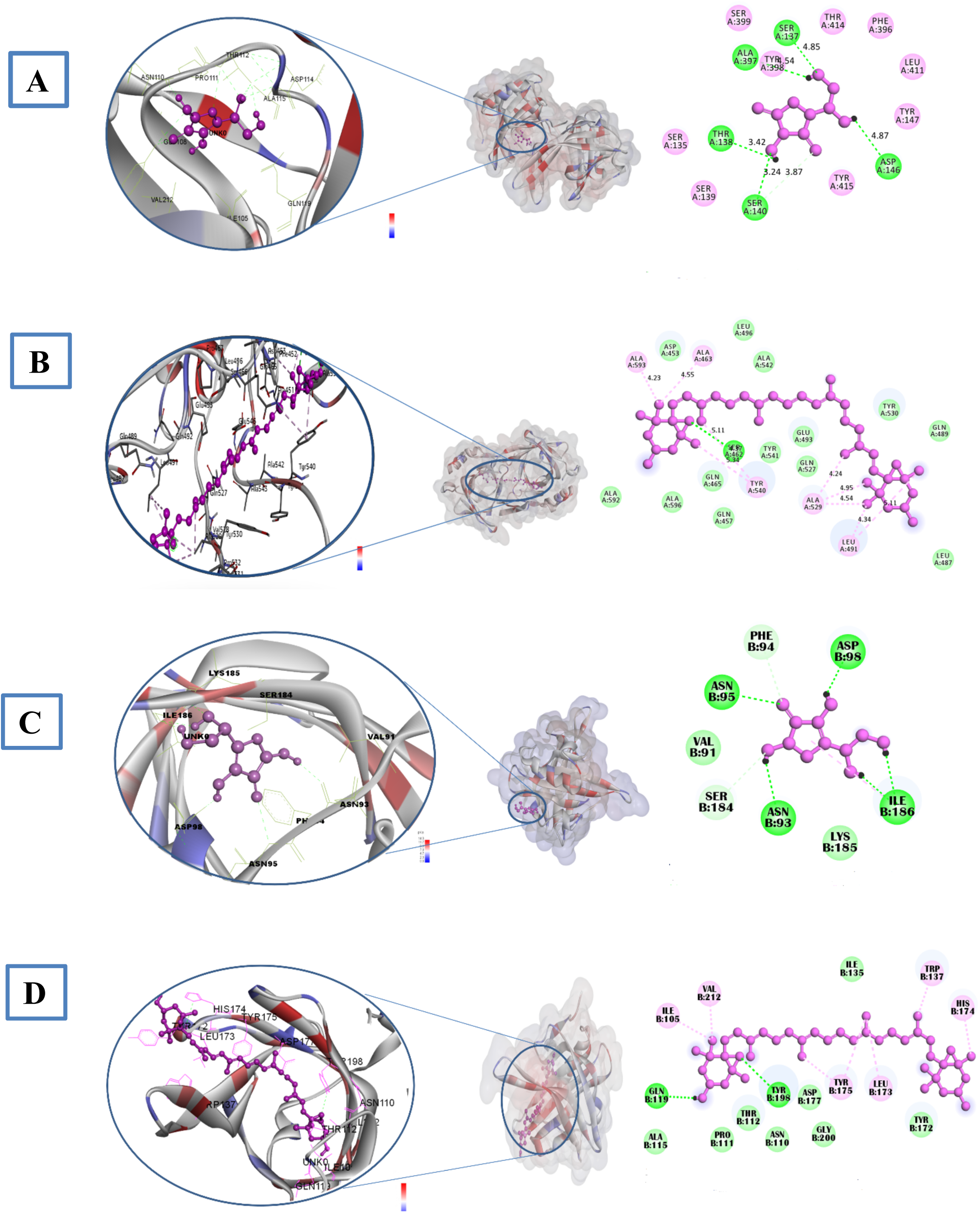
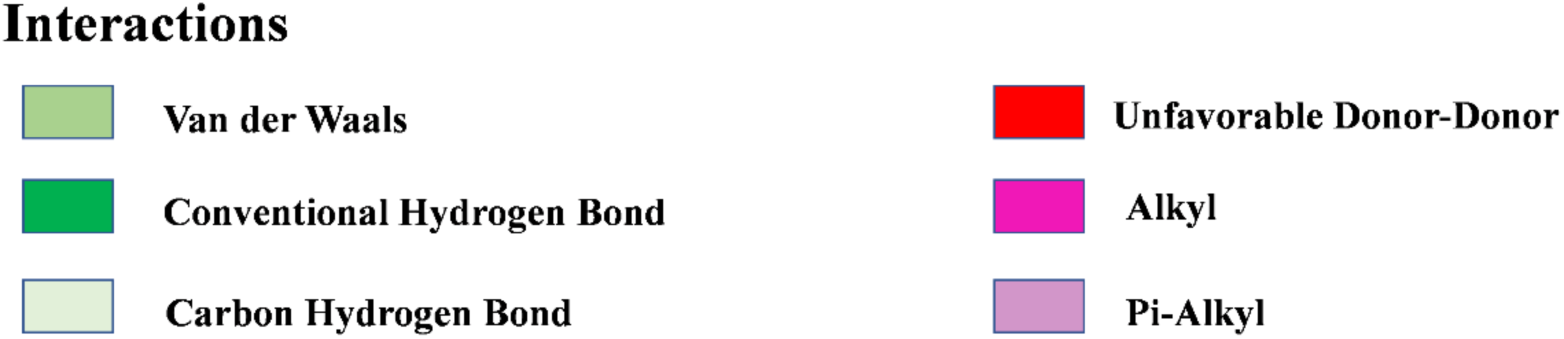
Interacting key residues and binding modes in (A) CVDE-ascorbic acid, (B) CVDE-violaxanthin, (C) VDE-ascorbic acid, and (D) VDE-violaxanthin complexes.

### 3.4. Salt-bridge Analysis

Salt-bridge plays a pivotal role in the thermo-stability of enzymes aiding to deduce different plausible reaction mechanisms[Charbonneau and Beauregard, 2013]. Table S4 provides details of salt-bridges and pK_a_ of residues in the enzyme complexes. While the initial configurations of the complexes possess a very few intramolecular salt-bridges, there is a significant increase in number after the MD run. All over the CVDE complexes retain a substantial number of salt-bridges compared to that of VDE. In reference to the initial configurations identified with 3, 4, 13, and 57 salt-bridges, the optimized complexes after MD have 37, 50, 232, and 245 salt-bridges, respectively, for VDE-ascorbic acid, VDE-violaxanthin, CVDE-ascorbic acid, and CVDE-violaxanthin. Moreover, CVDE-violaxanthin and CVDE-ascorbic acid possess namely 103 and 87 new salt-bridges, which are formed at the side chains contributing to the self-stability of the protein[Charbonneau and Beauregard, 2013; Meuzelaar et al., 2016; Pace et al., 2009]. There is no intermolecular salt-bridge seen between the protein enzymes and substrates whatsoever. Table S3 reports salt-bridges which sustain before and after the MD simulation. pK_a_ calculations across the residues demonstrate that a lower average pK_a_ value is crucial for having a greater number of salt-bridges in the complexes (see Table S4). This implies that lower the difference (Δ) of the average pK_a_ between residues with and without a salt-bridge, chances of having more salt-bridges in the complexes are higher and vice-versa.

Taken into account the total number of acidic and basic residues present in VDE and CVDE, which are actually responsible for the salt-bridge formation, the number of salt-bridges formed per residue is larger for the violaxanthin complexes than that of ascorbic acid. Again, this number is larger for CVDE than VDE. As there exists a larger average no. of salt-bridges in CVDE or the violaxanthin complexes, consequently the protein-protein interaction is more there and the scope of interaction with the substrate(s) is less resulting in a weaker bonding between the corresponding enzymes and the substrates, which justifies the similar trend of binding free energies reported in Table 2.

**Table 2:**
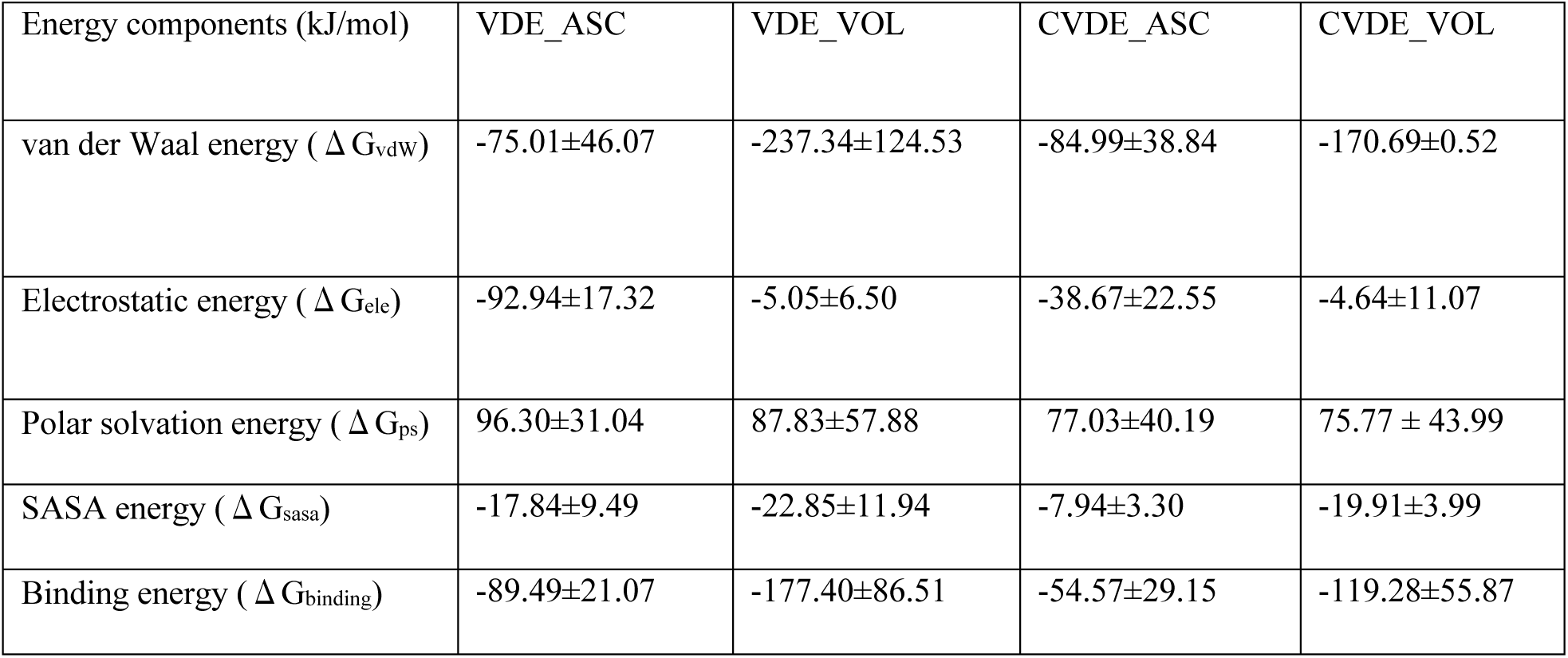
Binding free energies of the CVDE and VDE Complexes.

| Energy components (kJ/mol) | VDE_ASC | VDE_VOL | CVDE_ASC | CVDE_VOL |
| --- | --- | --- | --- | --- |
| van der Waal energy ( $\Delta G_{vdw}$ ) | -75.01±46.07 | -237.34±124.53 | -84.99±38.84 | -170.69±0.52 |
| Electrostatic energy ( $\Delta G_{ele}$ ) | -92.94±17.32 | -5.05±6.50 | -38.67±22.55 | -4.64±11.07 |
| Polar solvation energy ( $\Delta G_{ps}$ ) | 96.30±31.04 | 87.83±57.88 | 77.03±40.19 | 75.77 ± 43.99 |
| SASA energy ( $\Delta G_{sasa}$ ) | -17.84±9.49 | -22.85±11.94 | -7.94±3.30 | -19.91±3.99 |
| Binding energy ( $\Delta G_{binding}$ ) | -89.49±21.07 | -177.40±86.51 | -54.57±29.15 | -119.28±55.87 |

### 3.5. MD Trajectory Analysis

From the MD simulations, the intrinsic dynamical stability of each complex was studied by the root-mean-square deviation (RMSD) of the backbone, radius of gyration (R_g_), solvent accessible surface area (SASA) and root-mean-square fluctuation (RMSF) of the C_α_ atom as a function of simulation time, which are shown in Figure 2. RMSD measures the difference between the final and initial positions of the backbone of a protein. By means of deviations plotted over the course of simulation, the stability of a protein-ligand complex relative to its preliminary state can be easily assessed. A smaller deviation indicates the greater stability of protein-ligand complexes. Of the VDE- and CVDE-ascorbic acid complexes, the RMSD plot of VDE is smooth and nearly flat with an average backbone deviation of 0.45 Å in comparison to 0.63 Å of the latter where fluctuations as well as an increasing trend of RMSD are obvious, see Figure 2(i) A & A1. Exhibiting quite similar trends between VDE- and CVDE-violaxanthin complexes as of the ascorbic acid complexes, the average backbone deviations are, respectively, 0.41 Å and 0.76 Å, which are the highest and the least value of RMSD among all and so does their stability (Figure 2(ii) A & A1).

**Figure 2:**
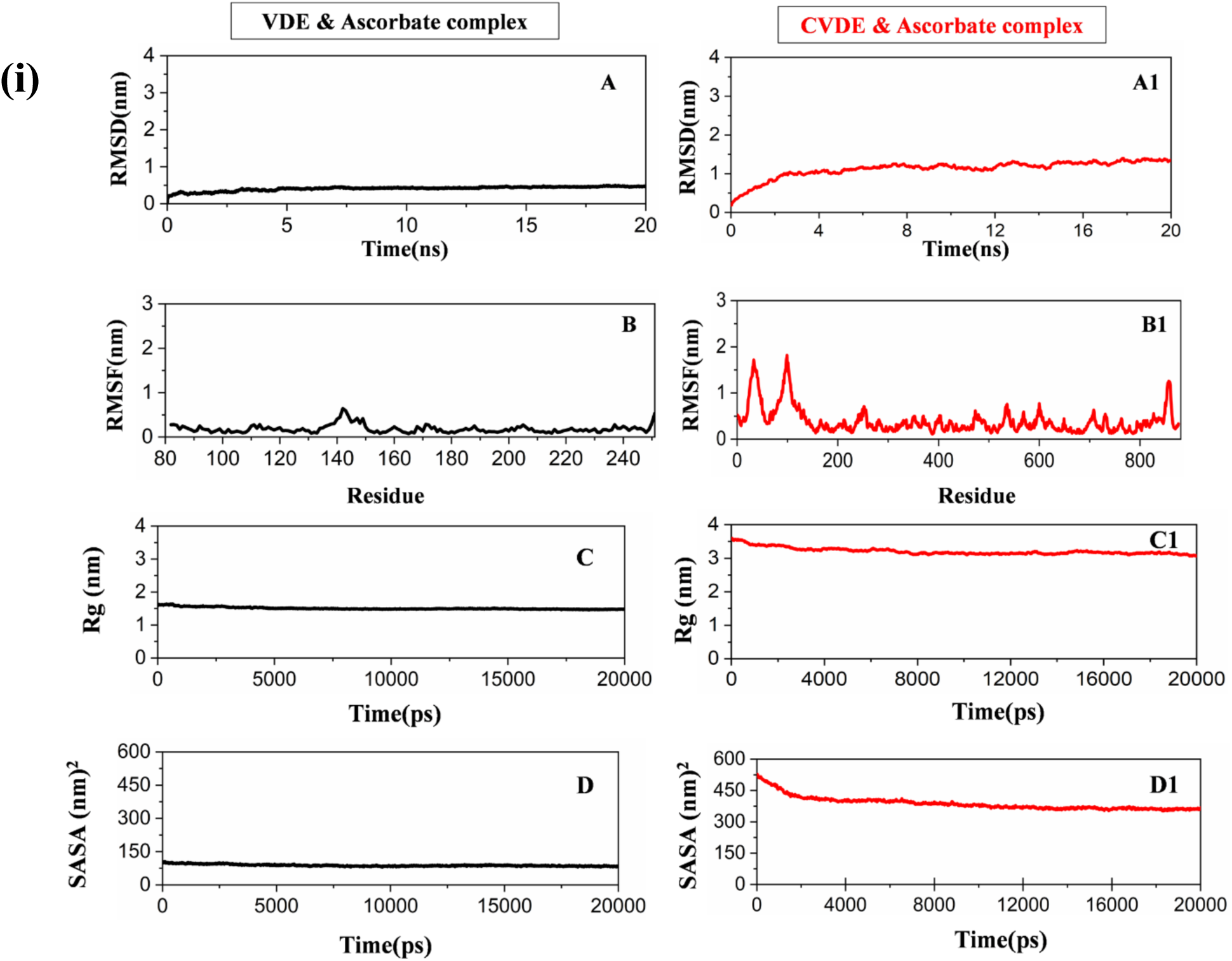
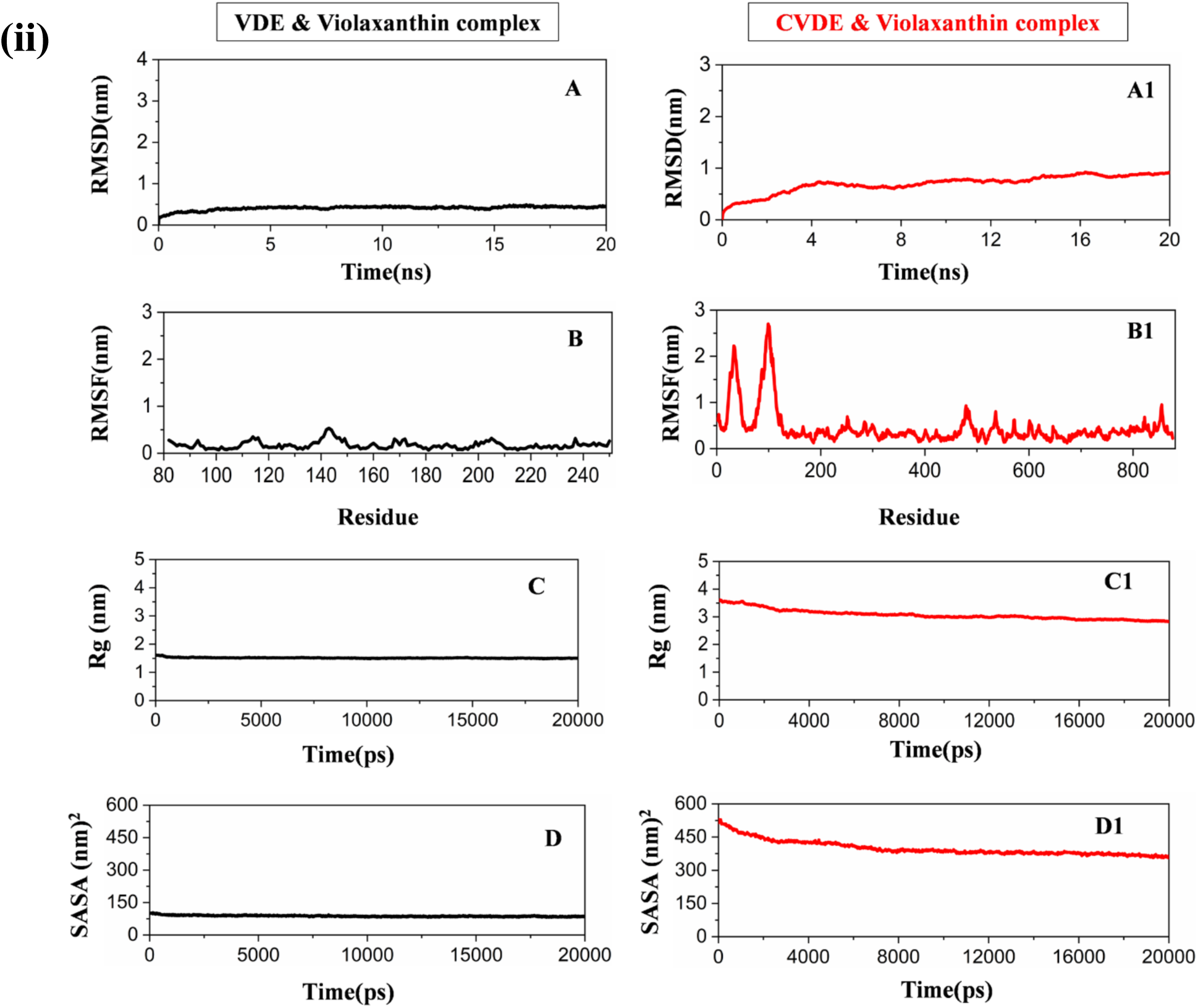
Plots of the root-mean-square deviation (RMSD) (A & A1), root-mean-square fluctuation (RMSF) (B & B1), radius of gyration (R_g_) (C & C1), and solvent accessible surface area (SASA) (D & D1) of ascorbic acid (i) and violaxanthin (ii) complexes of VDE (black) and CVDE (red).

The radius of gyration (R_g_) plots about a common center of mass are shown in C & C1 of Figure 2(i) & (ii), with the CVDE complexes having larger values than the VDE complexes. In the VDE complexes, the average R_g_ values are 1.56 Å and 1.59 Å, respectively, for ascorbic acid and violaxanthin, whereas 2.64 Å and 3.76 Å are the same for the CVDE complexes. Hence, in terms of forming a more compact complex, VDE is better than CVDE.

Otherwise, this becomes transparent in the plots of the solvent accessible surface area (SASA) of the proteins, D & D1 of Figure 2(i) & (ii), where the complexes of VDE have smaller surface areas than that of CVDE with the average values, respectively: 77.9 and 379 Å^2^ (for violaxanthin); 85.6 and 363 Å^2^ (for ascorbic acid). Towards ligand binding strength and the stability of the complexes, the RMSF plots (B & B1 of Figure 2(i) & (ii)) for the C_α_ atom and the average values are also quite insightful. Compared to VDE, the plots for CVDE contain more noises and larger RMSF values, which indicate relatively a weak binding between CVDE and the substrates. As the average RMSF increases in the order for VDE: VDE-violaxanthin (0.25 nm) < VDE-ascorbic acid (0.32 nm) and in case of CVDE: CVDE-violaxanthin (0.72 nm) < CVDE-ascorbic acid (0.87 nm), the binding strength and hence the stability of the protein-substrate complexes should decline in the same sequence. Worthwhile to mention that a few regions including the variable loops show greater fluctuations in their C_α_ atoms in all the systems indicating more flexibility of these residues. In both the complexes of CVDE, a significant fluctuation occurs at aa residues 50 to 60 and 90 to 100, with the highest peak located at the 100^th^ aa residue. Some extra fluctuations after 860^th^ residue also contribute to the RMSF of CVDE-violaxanthin.

### 3.6. Intermolecular H-bond Analysis

H-bond is one of the major driving interactions contributing to the stability of the complexes. Using gmx hbond, h-bonds formed over the simulation period were analyzed in all the complexes and are presented in Figure 3. Compared to the ascorbic acid complexes, the violaxanthin complexes manifest a consistent pattern of h-bond formation. Among all, VDE-ascorbic acid has by far the largest number of h-bonds (average: 3.91) and the corresponding CVDE complex has the lowest number (average: 1.33). On the contrary, the violaxanthin complexes of VDE and CVDE have a consistent pattern over time with the average numbers of h-bonds being comparable, 1.95 and 2.21, respectively. Considering the overall aspect of dynamical variations of the intermolecular h-bond and binding affinity, it is immediately evident that CVDE and VDE form more stable complexes with violaxanthin than with ascorbic acid.

**Figure 3:**
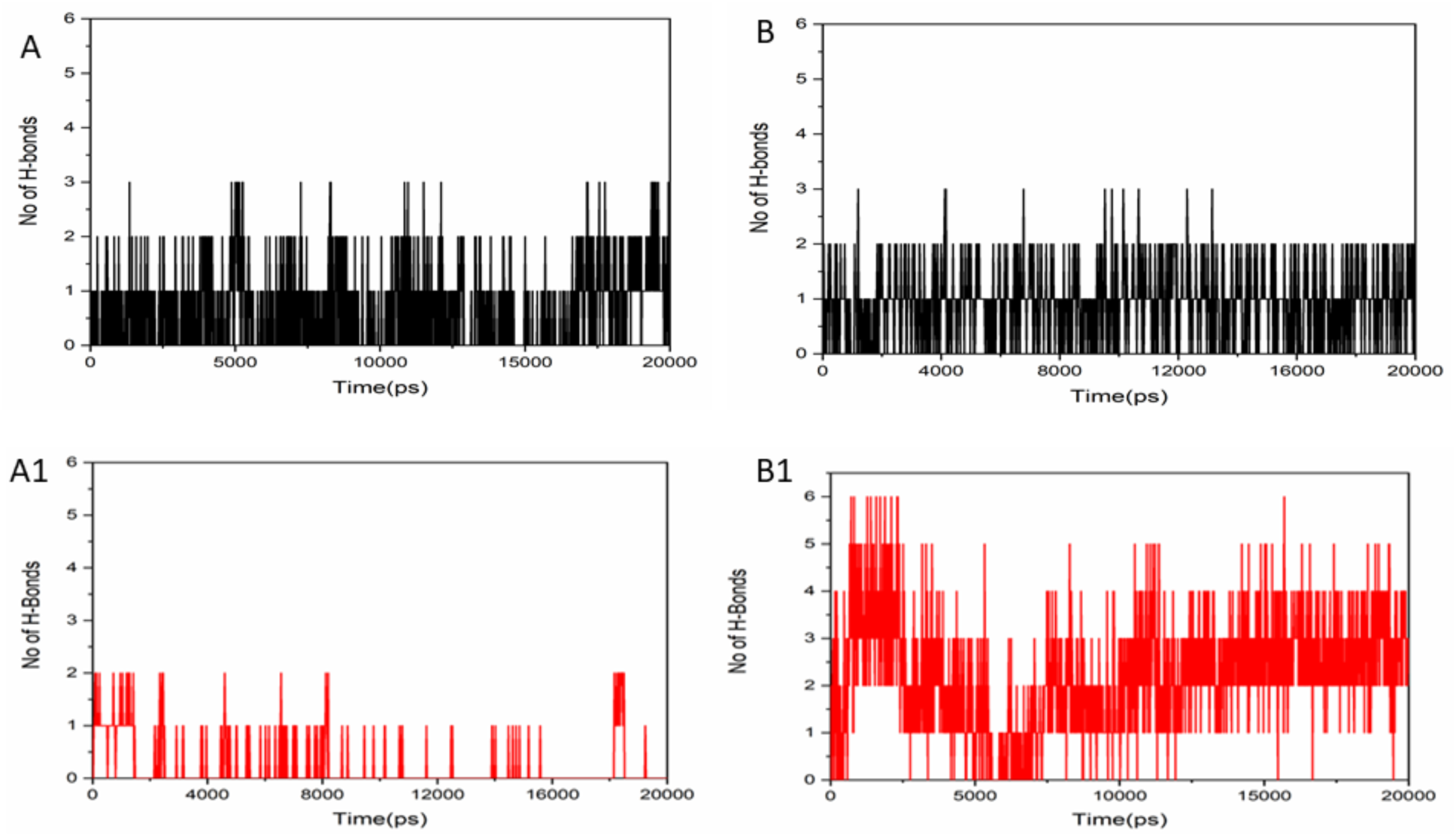
Plots of the average number of intermolecular h-bonds in the ascorbic acid (red) and violaxanthin (black) complexes of (A1) CVDE and (B1) VDE.

### 3.7. Secondary Structure Analysis

With gmx DSSP tool of GROMACS, secondary structure analysis of all the complexes was performed. This study helps us to examine the secondary structure of proteins, which contains helices, beta-sheets and loops, and associated internal motions over time (see Figure 4). In both the complexes of CVDE, formation of coil is predominant, around 50% residues which are part of this don’t play any role in structure formation. Around 20% residues not forming any h-bonds are present, which also don’t help in structure build-up. Residues capable of exhibiting h-bond interactions i.e. alpha –helices account for another 20%. In passing, the presence of 5-helices and turns gives rise to a large number of h-bond formations in VDE-violaxanthin compared to VDE-ascorbic acid. Due to more parallel and antiparallel backbone h-bonds present in the beta-sheet, CVDE-violaxanthin has the more stable secondary structure than CVDE-ascorbic acid. While beta-sheets present at large proportion in both the complexes of VDE, most of the residues participate in structure formation. Interestingly, the maximum number of h-bonds shown by VDE-ascorbic acid arises from prevailing structural characteristics of different types of motifs helping in secondary structure formation such as alpha-helices, beta-sheet, beta-bridge, and turns. As most of the beta-bridge residues help in beta-sheet formation in CVDE-violaxanthin than CVDE-ascorbic acid, the latter exhibits inferiority in the formation of alpha-helices due to the lack of rigidity and stability.

**Figure 4:**
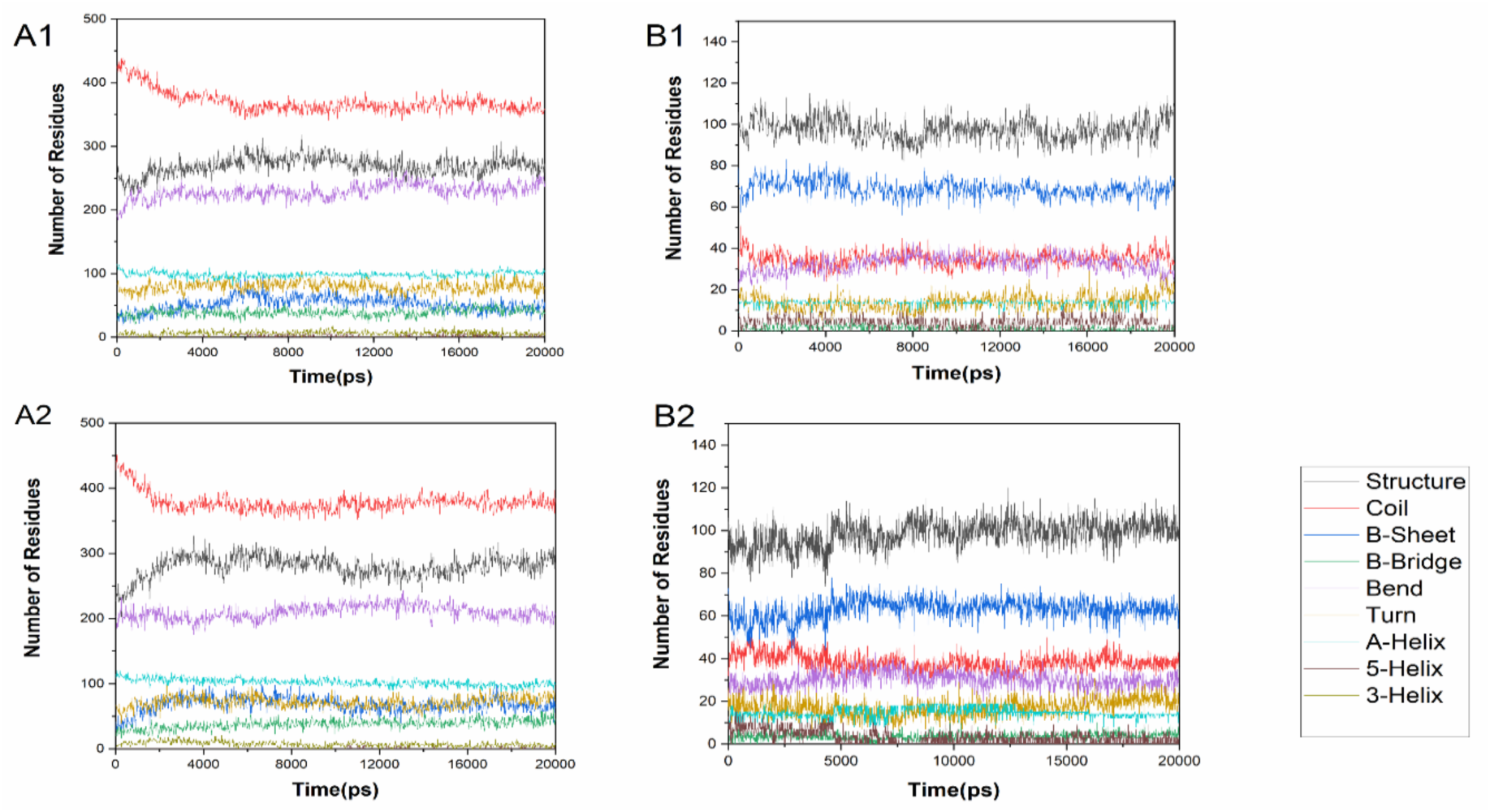
Secondary structure analysis of the CVDE and VDE complexes: (A1) VDE-ascorbic acid, (A2) VDE-violaxanthin, (B1) CVDE-ascorbic acid, and (B2) CVDE-violaxanthin.

### 3.8. Binding Free Energy Calculation

The binding free energy and its various components obtained from MM/PBSA [Kumari et al., 2014; Remko and von der Lieth, 2006] calculations of the CVDE and VDE complexes are listed in Table 5. Free energy calculations indicate that both CVDE and VDE with violaxanthin possess the highest negative binding free energy of -119.28 ± 55.87 kJ mol, -177.40 ± 86.51 kJ mol, respectively, as compared to the corresponding ascorbic acid systems. In case of all the enzymes, the polar solvation (ps) energy is positive, against the binding with ascorbic acid and violaxanthin; while the rest such as the van der Waals (vdW), electrostatic (ele), and solvent accessible surface area (sasa) energies are negative favoring binding. Among the favorable interactions, ele and sasa are the least contributor, respectively, for the violaxanthin and ascorbic acid acid complexes. Whereas, the vdW interaction has the largest contribution to binding almost for all. The high negative value of vdW energy suggests that a hydrophobic interaction substantially interplays between VDE and violaxanthin, CVDE and violaxanthin/ascorbic acid. Based on the binding free energy analysis, it can be demonstrated that both CVDE and VDE systems with violaxanthin appear to exhibit the most intrinsic dynamical stability than with ascorbic acid. This is in consistent with the earlier prediction based on the binding energies reported in Table 2. These findings explain as to why VDE and violaxanthin taking part in many catalytic processes reported in the literature. In the experimental work of Meireles *et al*.^31^, the ascorbate deficiency does not affect the de-epoxidation process in CVDE, suggesting that ascorbate is not required for this process. The lower negative binding energy/free energy of CVDE-ascorbic acid here indeed confirms this experimental finding, undermining the role of ascorbic acid in de-epoxidation for CVDE compared to violaxanthin.

### 3.9. PCA

In PCA, essential dynamics (ED) was employed to represent the directions of principal motions by a set of eigenvectors (EV) called principal or essential modes. Each MD trajectory was projected onto the phase space to yield a spectrum of EVs, which depict the vectorial representation of each single components in motion. Each EV holds an eigenvalue that describes the energetic contribution of each component to the motion (Figures 5A, C, E, and G). It is observed that 75-77% of the backbone motion is covered by the first 20 EVs where an exponentially decaying curve of eigenvalues is obtained against the EVs (Figure 5). The differential scattering of atoms in contour-like plots specifies the occurrence of conformational changes in the complexes in agreement with other MD analyses. Figures 5B, D, F, and H reveal a higher subspace dimension in both the complexes of CVDE. In contrast to CVDE, the VDE systems cover the least subspace and show the least variations while the first two PCs were taken into consideration. In the 2-D projection plots of trajectories, the CVDE systems show higher trace values of the covariance matrix than the VDE systems. In the overall PCA analysis, we conclude that the CVDE systems have more flexibility than the VDE systems in which the VDE-violaxanthin complex shows the least conformational changes due to decreasing collective motions than other systems. This result is in congruent with the stability of the VDE-violaxanthin complex in the xanthophyll cycle during de-epoxidation chemical process. Emphatically, the PCA and RMSF analyses elicit the same characteristics about structural fluctuations of the protein-substrate complexes.

**Figure 5:**
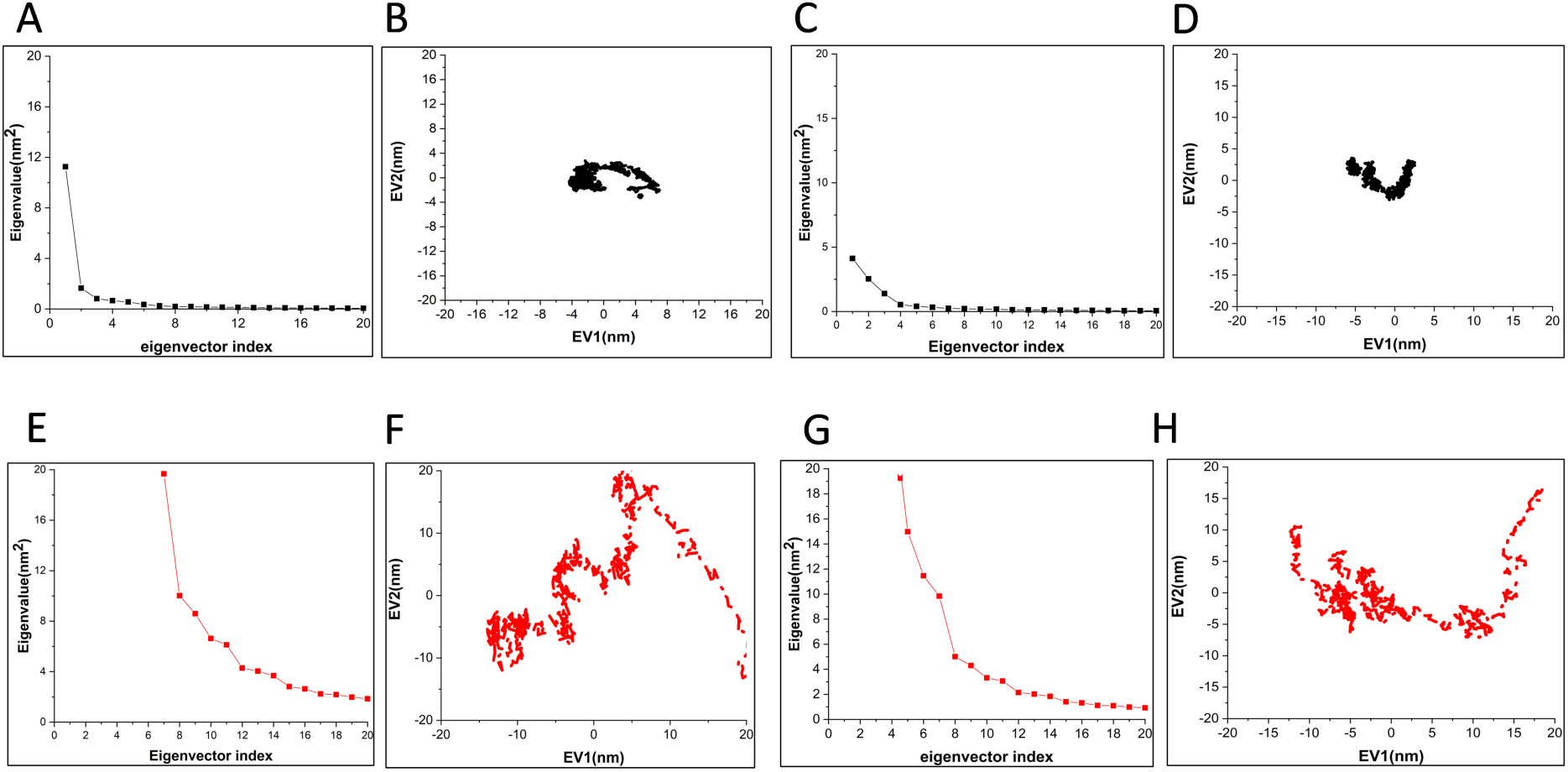
Principal component analysis of the CVDE and VDE complexes: plot of eigen values versus the corresponding eigen vector indices coming from the C_α_ covariance matrix during MD simulations (A, C, E, and G) and the 2-D projection plot of the first two principal eigenvectors (B, D, F, and H). [Black: VDE complexes with both ascorbic acid (A & B) and violaxanthin (C & D); Red: CVDE complexes with both ascorbic acid (E & F) and violaxanthin (G & H)].

## 4. Conclusion

Violaxanthin de-epoxidase (VDE) plays a pivotal role in xanthophyll cycle and keeps photosynthesis process intact by protecting from reactive oxygen species generated during the process. Very recently, a variant of VDE (called Chlamydomonas CVDE) present in algae is discovered, which functionally resembles with VDE enzyme but was not explored. To understand whether there exist any structural and functional similarities, a comparative study is performed between VDE and CVDE shedding light on the primary steps of the xanthophyll (photosynthesis) cycle. Hence, phylogenetic profile, Bioinformatics analyses for structure determination and modeling, molecular dynamics studies (to check binding stability, fluctuations, energetics, etc.) of the enzyme complexes with the same substrates are performed. It is found that CVDE is an ancestor of VDE. Like VDE, CVDE has a catalytic domain even larger than that of VDE, thus can exhibit catalytic properties. From the essential dynamics (ED), RMSD, R_g_, and intermolecular h-bonds, CVDE displays higher conformational changes compared to VDE, which may attribute to its larger structure having more residues than VDE. Evaluation and decomposition of binding free energies by MM/PBSA demonstrate that the van der Waals and electrostatic interactions majorly contribute to binding with the substrates for both enzymes. Along with the binding energies, the free energy analysis enlightens that both CVDE and VDE have equal affinity to violaxanthin, while VDE binds to ascorbic acid stronger than CVDE. We found out that the experimental result of Meireles *et al*. is in congruent with our study, indicating that the de-epoxidation process in CVDE requires different reductants than ascorbic acid. Overall, the study discerns that CVDE might have similar roles as VDE in photosynthesis/photoprotection in the presence of violaxanthin and trivial roles with ascorbic acid. Taken everything into consideration, this work presents new insights into structural details and binding mechanisms of CVDE that were unknown till date as compared to widely studied VDE, which may incite more theoretical and experimental studies in this direction as far as photosynthesis or the xanthophyll cycle is concerned.

## Conflict of interest

The authors declare no conflict of interest and all the authors have approved this article for publication.

## Acknowledgements

The authors acknowledge IISER Berhampur for computational support. P.S.S.G also sincerely acknowledges IISER Berhampur for providing him the Institute Postdoc Fellowship to carry out this work.

## Notes

### Competing Interest Statement

The authors have declared no competing interest.

