## Supplementary Materials for "Insights into the Binding Mechanism of Ascorbic Acid and Violaxanthin with Violaxanthin De-Epoxidase (VDE) and Chlorophycean Violaxanthin De-Epoxidase (CVDE) Enzymes: Docking, Molecular Dynamics, and Free Energy Analysis"

### SUPPLEMENTARY MATERIAL

**Table of contents:**

**Figure S1:** Multiple Sequence alignment of CVDE: target-template alignment of CVDE with the close structural homolog.

**Figure S2:** Secondary structural information prediction of CVDE by PSIPRED server.

**Figure S3:** Structural overview of CVDE 3D model: (A) 3-D architecture of zCDKL5 (The protein is shown as solid ribbon representation with α-helices, β-sheets, and coils, (B) Ramachandran plot of the modeled domains, and (C) topology of the CVDE 3-D model

**Table S1:** Comparative analysis among various model validation approaches between the homolog modeled CVDE and VDE (PDB ID: 3QCR).

**Table S2:** Calculated electronic properties of ascorbic acid and violaxanthin using DFT/B3LYP/6-31+G (d, p).

**Table S3:** Existing salt-bridges in the complexes of (a) VDE-ascorbic acid, (b) VDE-violaxanthin, (c) CVDE-ascorbic acid, and (d) CVDE-violaxanthin before and after the MD simulation.

**Table S4**: Number of salt-bridges present in each complex, corresponding average $pK_a$ over all residues, over salt-bridge forming (x) and not forming (y) residues, and their $pK_a$ difference ( Δ = y-x).

### SUPPLEMENTARY FIGURES

**Figure S1: Multiple Sequence alignment of CVDE: target-template alignment of CVDE with the close structural homolog.**

```
                      260       270       280       290       300       310
9synCruP       PAHDGR..TTYLFTYLDAHPQRF.DLAFLLEEYLKLLPDYQQVDLAALDFQRFLFGFFPS
5CyaCruP       PAKDGR..TTYLFTYVDAHPDRI.SLTDLTQEYLRLLPEYQNIDLGAIAFKRFLFGFFPS
9SynCruA       PGQGDEM.TIYLFHYHEVNPENPGSLLEMYEDFFSILPEYRRCDMAQLTFEKATFGYIPG
5CyaCruA       PAERDEL.TIYLFHYHQVHPDNPGSLLEMYEDFFNILPEYRRCDMDKLVWKKPTFGYIPG
1CrCVDE        PAEGGQARTTYMFAYSDANPDRP.SFEALLDTYPQMLPEYQGIPLDQLKFKRVLFGGFPC
3CruCruP       ALEPS.HVLKAEAGRIEVWDHHAPQLRCSGVSFVRYIIESKGLYLPSFFSTARLSFVAKG
3CruCruA       TANPANHVIQAEYPDYYFRITNSEHMTDLKEKFKRMCXKSM...IRKRHMHLTEEFLKDN
consensus>50   pae..e..t.ylf.y.d..pd...sl.dl.ee%..llpey...dl..l.f...lfgf.p.


                      320       330       340       350       360
9synCruP       YRR...SPLHYPWDRILPIGDSSGGQSPVSFGGFGAMLRHLERLTNGLDDALTQDCCDRQ
5CyaCruP       YSN...SPLKMPWQRILAIGDSSGSQSPVSFGGFGSMMRHLPRLNEAINEALKTDSLSPK
9SynCruA       YFNVGAGDRQVAFDRLLAIGDAASLQSPLVFTGFGSLVRNLDRLTKLLDIALQKDLLDQQ
5CyaCruA       RFTVGENDREIATDRVMAIGDAASLQSPLVFTGFGSLVRNLGRLTSLLDKALKNDLLDGE
1CrCVDE        YSN...GPLAPAFDRVMQIGDASAAQSPLSPGGFGSMMAHLPRLARGLDQALQEDRLARP
3CruCruP       EGLMGRVVLCAETFQDSSVFQPSGGSPFGEGQGQGQQGQGQGHQGQGQG.QQGQQGQQGQ
3CruCruA       PDMCAYMAPSLDARQDIVVVEVPKLGKEAAVKAIKEWGQPKSKTTHVVFCTTSGVDMPGA
consensus>50   y.......l....dril.!g#.s..qsp..f.gfgsm.r.l.rlt..ld.al.qd.ld.q


               370       380       390        400       410       420
9synCruP       SLAQLQPYQPNLSVTWLFQKAMSVGVNQ.SCPPNQINDLLNAVFGVMAQLGEDTLNPFLQ
5CyaCruP       DLHLIQPYQPNISVTWLFQKTMSVGIDK.NYSPNQINDLMSGVFKVMDRMGDDVLKPFLQ
9SynCruA       NLSKIRAYQSNIAVTWLFSKGMMVPTGM.KLPPQRINAMLNTFFGLLADSSPEVAETFIK
5CyaCruA       SLNRIKAYQSNIAVTWLFSKGMMVPTHK.TLPPERVNAMLNTFFGLLADE.PDTAEIFIK
1CrCVDE        DLNWLHPYQPSLSASWLFQRSMSLAVGQVAYPPDCPHAPAYYAAAKEAKAAAAAAAVDRA
3CruCruP       QSQGQGFRDMHQKVEHIRTGDTIATHPGVAQWFYNDGNQPLVIVSVLDLASHQNQLD.RN
3CruCruA       DYQLTKLLGLRPSVKRLMMYQQGCFAGGTVLRLAKDLAENNRGARVLVVCSEITAVTFRG
consensus>50   dln....yq.n.svtwlf.k.m.v.......ppn.inalln..f.vla....d.a..f..


                  430              440       450       460       470
9synCruP       DVVQFQG.......LTKTLPRVNFKTVLPLLPHLGVGALADWLRHYLALGLYTSSYALSQ
5CyaCruP       DVVQFSG.......LAKTLPLVNPLLVLPLLPQVGLPVFADWFKHFTNLGLFTGLYPLAK
9SynCruA       DRTSWLMFNK....LALVAARQNPALLVWIWQMAGAKD...FI.RWVGAYFAFSFDAVLS
5CyaCruA       DRTDWFTFNR....LALKAARKNPALLLWIWEMAGTRD...IF.RWMGSYAKFTFDALKN
1CrCVDE        EGFDGLVSTAGERALSLQEAAMEAVEAVAARFAAGSADPADYF.HVEQEVPGAGSDRRTP
3CruCruP       PRPFYLAGN..........NPQGQVWIEGREQQPQKNILNGFTPEVLAKAFKIDVRTAQQ
3CruCruA       PSDTHL..D..........SLVGQALFSGGAAA...LIVGSDPDITAGEKPIFEMVSAA
consensus>50   d..q.l..n.....l.......n..l........g.........................


              480       490       500       510
9synCruP       RLPMGDSYQAKRRREAWQYGSGQDFHQAGLTEQD..........................
5CyaCruP       SFKSLES..TMNEQQLYYYHRYLDMWKYGAGKDN..........................
9SynCruA       LLLMGWLPQWLENSEAWLSEKYPSLWLSLLSLKYRLTVGT....................
5CyaCruA       LLFSRWFNPWLENQSKWLRESNPKLWFKLLTFGSRFVSQK....................
1CrCVDE        QLASGKAQPAPPKLKKKLFERDFRTAPEWQRLPYTHVNEILGTNFGVMGVLGDRVLKPFL
3CruCruP       LQNQQDNRGNIIRVQGPFSVIRPPLRSQRPQEEVNGLEETICSARCTDNLDDPSNADVYK
3CruCruA       QTILPDSDGAIDGHLREVGLTFHLLKDV.PGLISKNIEKSLDEAFKPLGISD.WNSLFWI
consensus>50   .l...d.....e..e.........l...................................
```

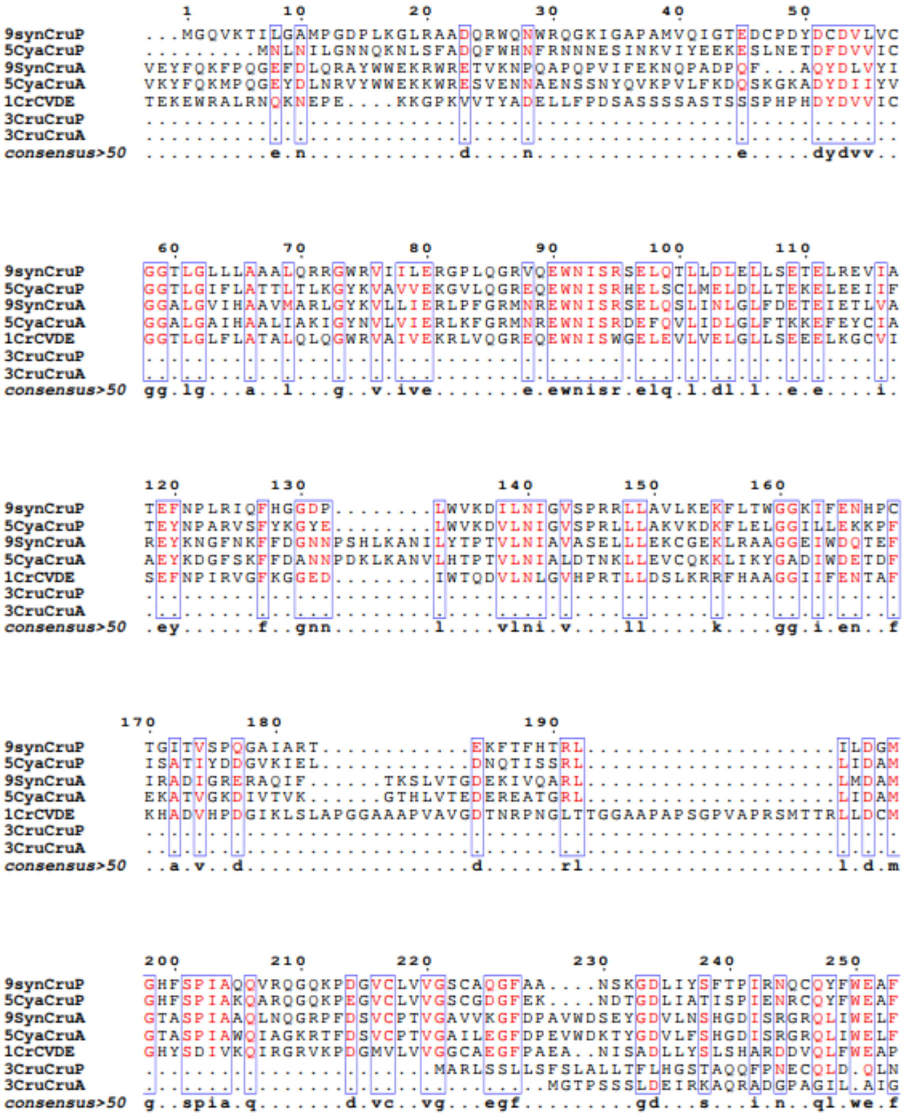

**Figure S2: Secondary structural information prediction of CVDE by PSIPRED Server.**

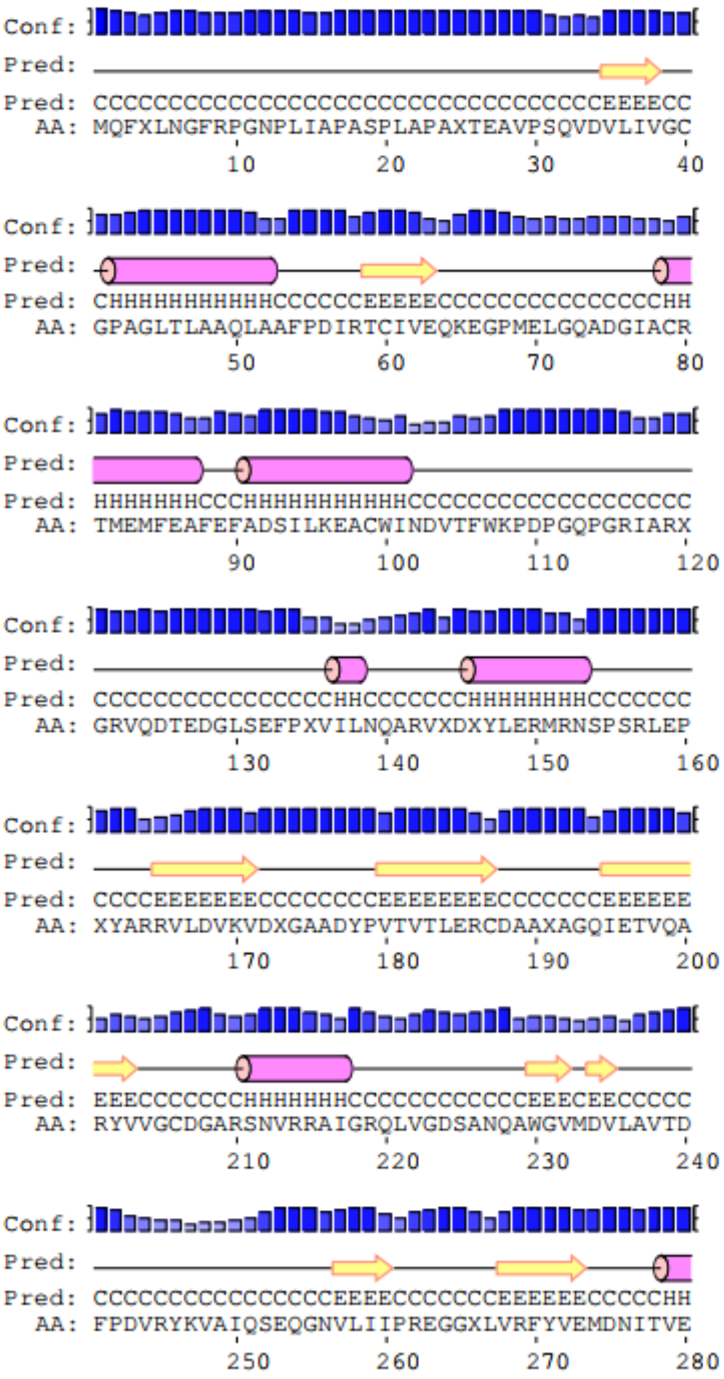

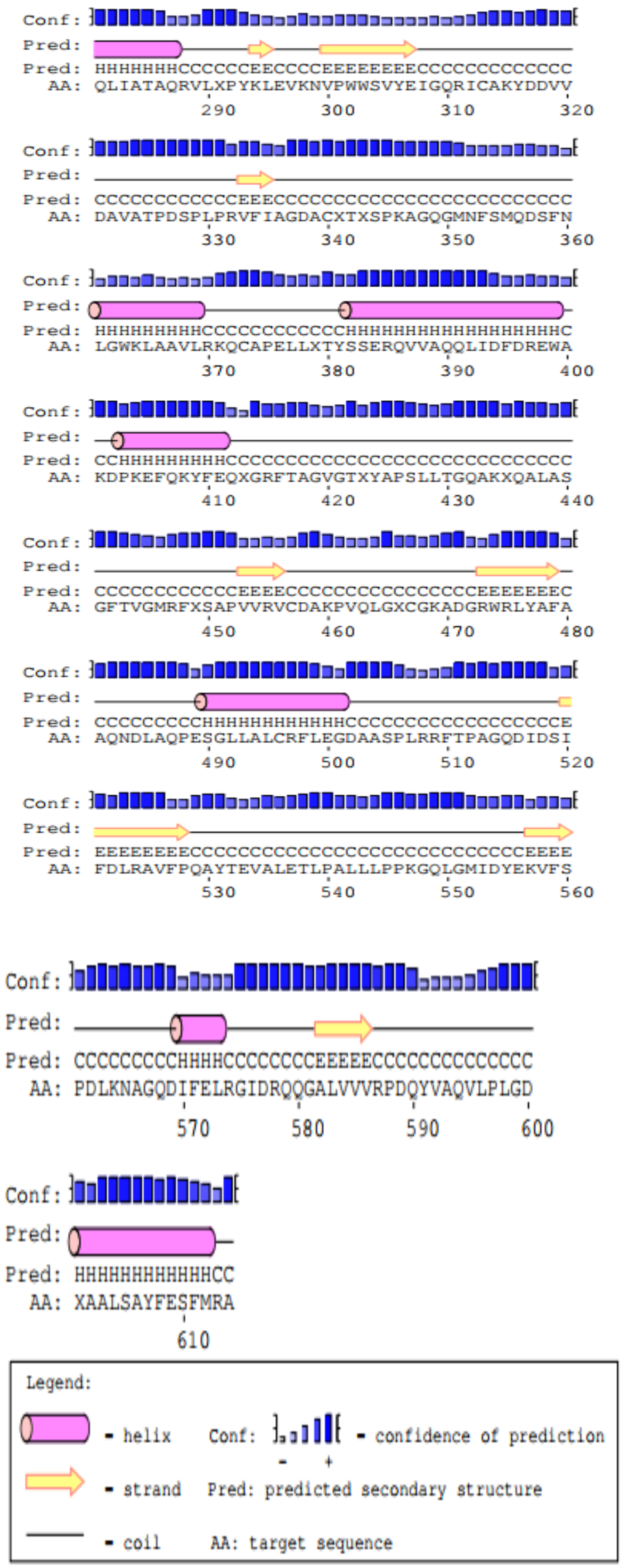
Conf:
Pred:
Pred: HHHHHHHCCCCCCEECCCCEEEEEEEEECCCCCCCCCCCC
AA: QLIATAQRVLXPYKLEVKNVPWWSVYEIGQRICAKYDDVV
290 300 310 320
Conf:
Pred:
Pred: CCCCCCCCCCCCEEECCCCCCCCCCCCCCCCCCCCCCCCC
AA: DAVATPDSPLPRVFIAGDACXTXSPKAGQGMNFSMQDSFN
330 340 350 360
Conf:
Pred:
Pred: HHHHHHHHHCCCCCCCCCCCHHHHHHHHHHHHHHHHHHHC
AA: LGWKLAAVLRKQCAPELLXTYSSERQVVAQQLIDFDREWA
370 380 390 400
Conf:
Pred:
Pred: CCHHHHHHHHHCCCCCCCCCCCCCCCCCCCCCCCCCCCCC
AA: KDPKEFQKYFEQXGRFTAGVGTXYAPSLLTGQAKXQALAS
410 420 430 440
Conf:
Pred:
Pred: CCCCCCCCCCCCEEEECCCCCCCCCCCCCCCCEEEEEEEC
AA: GFTVGMRFXSAPVVRVCDAKPVQLGXCGKADGRWRLYAFA
450 460 470 480
Conf:
Pred:
Pred: CCCCCCCCCHHHHHHHHHHHHCCCCCCCCCCCCCCCCCCE
AA: AQNDLAQPESGLLALCRFLEGDAASPLRRFTPAGQDIDSI
490 500 510 520
Conf:
Pred:
Pred: EEEEEEEECCCCCCCCCCCCCCCCCCCCCCCCCCCEEEE
AA: FDLRAVFPQAYTEVALETLPALLLPPKGQLGMIDYEKVFS
530 540 550 560
Conf:
Pred:
Pred: CCCCCCCCCHHHHCCCCCCCCEEEEECCCCCCCCCCCCCC
AA: PDLKNAGQDIFELRGIDRQQGALVVVRPDQYVAQVLPLGD
570 580 590 600
Conf:
Pred:
Pred: HHHHHHHHHHHHCC
AA: XAALSAYFESFMRA
610
Legend:
- helix
- strand
- coil
Conf: - confidence of prediction
- +
Pred: predicted secondary structure
AA: target sequence

**Figure S3: Structural overview of CVDE 3-D model: (A) 3-D architecture of zCDKL5. (The protein is shown as solid ribbon representation with α-helices, β-sheets, and coils), (B) Ramachandran plot of the modeled domains, and (C) topology of the CVDE 3-D model.**

**A**

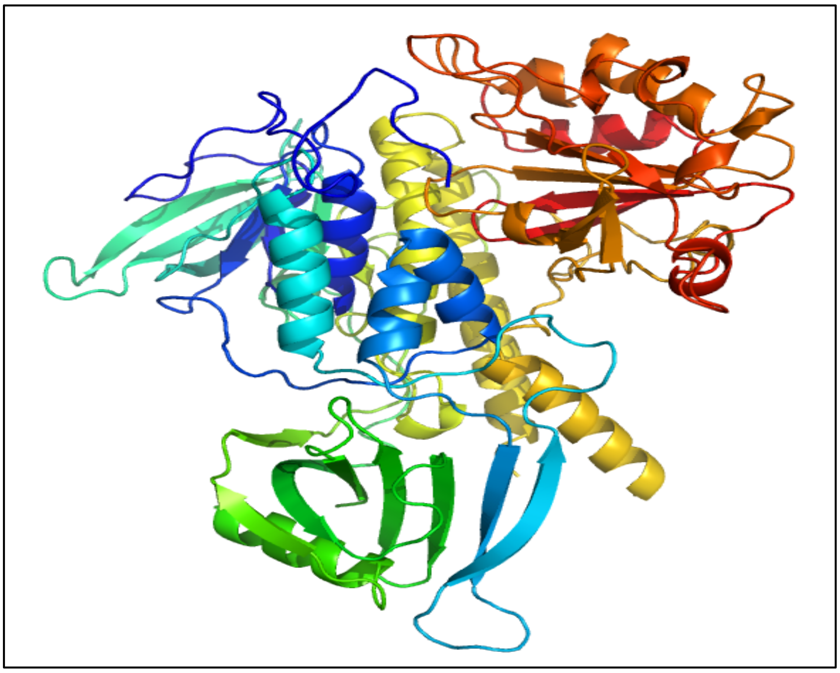

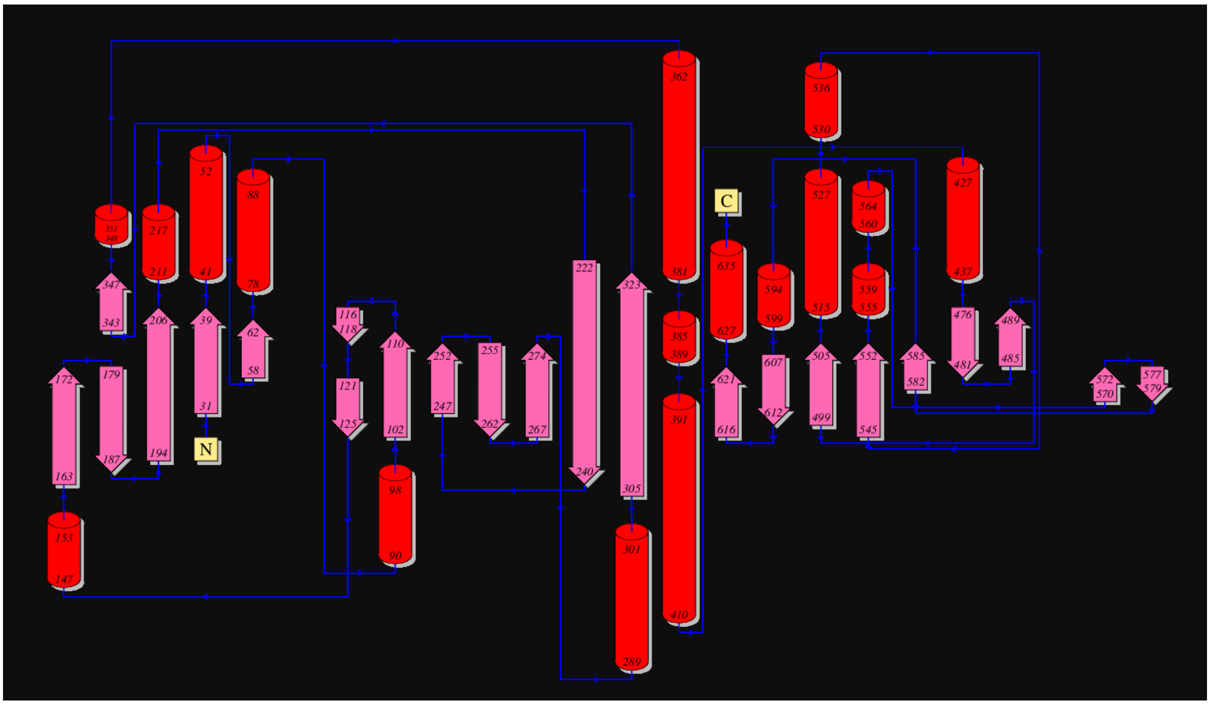

**B**

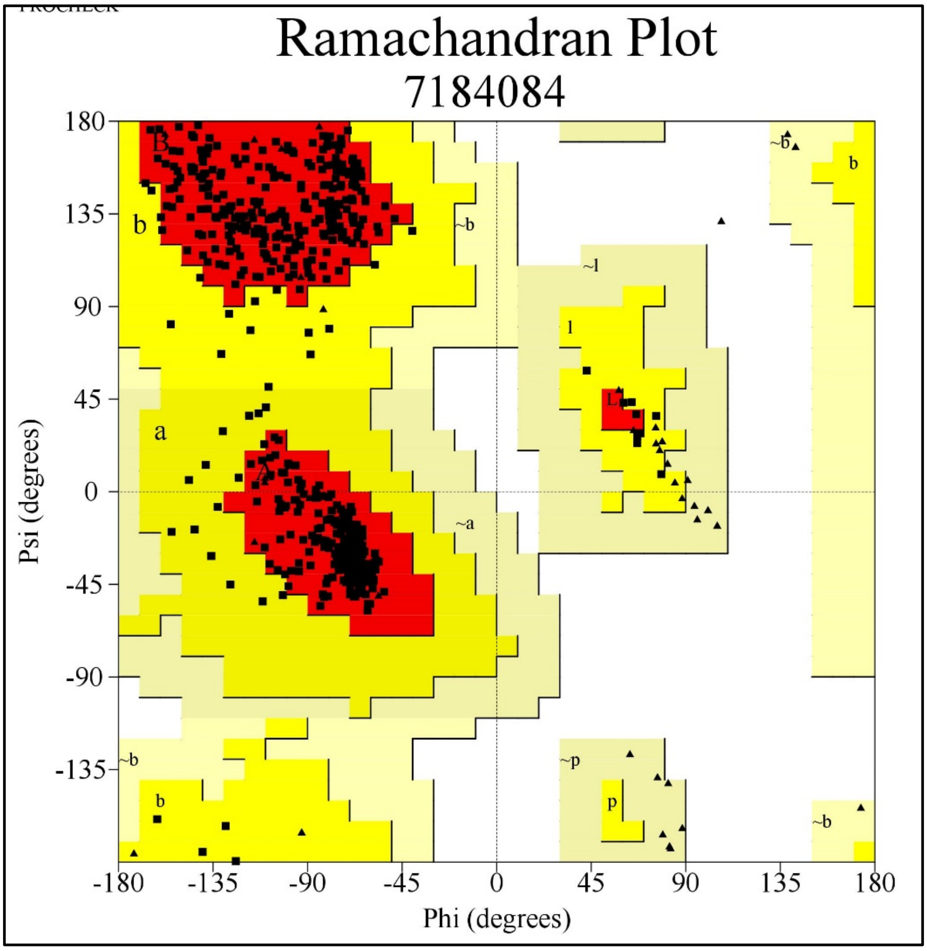

Ramachandran Plot
7184084
Psi (degrees)
Phi (degrees)
180
135
90
45
0
-45
-90
-135
-180
-135
-90
-45
0
45
90
135
180


**C**

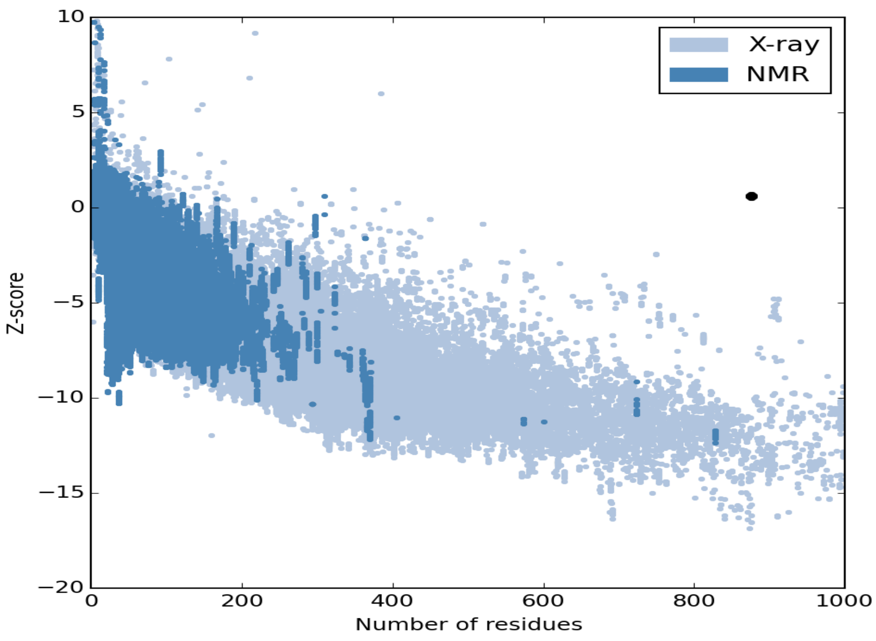

X-ray
NMR
Z-score
Number of residues

### SUPPLEMENTARY TABLES

**Table S1: Comparative analysis among various model validation approaches between the homolog modeled CVDE and VDE (PDB ID: 3QCR).**

| **Model Validation** | **Parameter** | **CVDE** | **VDE (PDB ID-3CQR)** |
|---|---|---|---|
| PROCHECK | PROCHECK Most favored regions (%) | 91.20 | 95.40 |
| | Additionally, allowed regions (%) | 6.20 | 4.60 |
| | Generously allowed regions (%) | 8.20 | 0.00 |
| | Disallowed region, % | 0.00 | 0.00 |
| Verify3D | Verify3D (Averaged 3D-1D score > 0.2 | 88.79 | 91.60 |
| ERRAT | Overall G-factor | 0.02 | 0.02 |
| | ERRAT Overall quality | 91.86 | 93.65 |
| ProSA | ProSA Z-Score | -4.54 | -5.77 |
| ProQ | ProQ LG score | 3.40 | 4.40 |
| MolProbity | MaxSub | 0.51 | 0.59 |
| | MolProbity $C_\beta$ deviations > 0.25 Å | 0.89 | 40.25 |
| | MolProbity Residue with bad bonds, % | 0.91 | 0.20 |
| | MolProbity Residue with bad angles, % | 0.51 | 0.18 |

**Table S2:** **Calculated electronic properties of ascorbic acid and violaxanthin using DFT/B3LYP/6-31+G(d, p).**

| **Electronic Descriptors (au)** | **Ascorbic Acid** | **Violaxanthin** |
|---|---|---|
| Total Energy (Hartree) | -676.59 | -1836.34 |
| Dipole Moment (Debye) | 4.04 | 2.52 |
| HOMO Energy (eV) | -0.35 | -0.24 |
| LUMO Energy (eV) | 0 | 0.05 |
| Band Gap (eV) | 0.35 | 0.29 |
| Ionization Potential (IP) (eV) | 0.35 | 0.24 |
| Electron Affinity (EA) (eV) | 0 | -0.05 |
| Chemical Potential ($\mu$) (eV) | -0.17 | -0.10 |
| Electronegativity ($\chi$) (eV) | 0.17 | 0.10 |
| Chemical Hardness ($\eta$) (eV) | 0.17 | 0.15 |
| Chemical Softness ($\sigma = 1/\eta$) (eV) | 5.75 | 6.84 |
| Electrophilicity Index ($\omega$) (eV) | 0.09 | 0.03 |

**Table S3: Existing salt-bridges in the complexes of (a) VDE-ascorbic acid, (b) VDE-violaxanthin, (c) CVDE-ascorbic acid, and (d) CVDE-violaxanthin before and after the MD simulation.**

| Residue group 1 (atomic no & residue) | Residue group 2 (atomic no & residue) | Distance in Å (before MD) | Distance in Å (after MD) |
|---|---|---|---|
| VDE-Ascorbate | | | |
| LYS 130 | GLU 126 | 2.74 | 3.11 |
| LYS 154 | GLU 126 | 3.92 | 2.44 |
| VDE-Violaxanthin | | | |
| LYS 102 | GLU 122 | 3.43 | 2.92 |
| HIS 168 | ASP 178 | 2.57 | 3.36 |
| HIS 168 | ASP 178 | 3.79 | 3.70 |
| CVDE-Ascorbate | | | |
| ARG 52 | ASP 57 | 3.71 | 3.42 |
| LYS 99 | ASP 95 | 1.21 | 1.71 |
| LYS 99 | GLU 98 | 2.29 | 2.33 |
| LYS 223 | ASP 272 | 3.95 | 2.69 |
| HIS 239 | GLU 840 | 3.57 | 3.80 |
| ARG 338 | ASP 364 | 3.79 | 3.59 |
| ARG 434 | GLU 646 | 2.66 | 3.89 |
| ARG 495 | ASP 227 | 3.77 | 3.98 |
| ARG 704 | ASP 358 | 3.58 | 3.77 |
| CVDE-Violaxanthin | | | |
| LYS 99 | ASP 95 | 3.19 | 0.63 |
| LYS 223 | ASP 272 | 3.95 | 3.01 |
| HIS 239 | GLU 840 | 3.57 | 2.86 |
| ARG 338 | ASP 364 | 3.79 | 3.69 |

| | | | |
|---|---|---|---|
| ARG 391 | GLU 185 | 3.83 | 3.88 |
| ARG 495 | ASP 227 | 3.77 | 3.72 |
| ARG 704 | ASP 358 | 3.33 | 3.46 |

**Table S4**: **Number of salt-bridges present in each complex, corresponding average $pK_a$ over all residues, over salt-bridge forming (x) and not forming (y) residues, and their $pK_a$ difference ( Δ = y-x).**

| Complexes | Total Number of Salt-bridges | Total Number of Acidic and Basic Residues | Number of Salt-bridges per Residue | Average $pK_a$ over All Residues | Average ($pK_a$) over Residues | | Difference (Δ = y-x) |
|---|---|---|---|---|---|---|---|
| | | | | | with Salt-bridge (x) | without Salt-bridge (y) | |
| VDE–ascorbic acid | 37 | 35 | 1.06 | 6.72 | 6.45 | 6.98 | 0.53 |
| VDE–violaxanthin | 50 | 35 | 1.43 | 6.87 | 6.85 | 6.89 | 0.04 |
| CVDE–ascorbic acid | 232 | 163 | 1.42 | 8.18 | 7.38 | 8.97 | 1.59 |
| CVDE–violaxanthin | 245 | 163 | 1.5 | 8.04 | 8.03 | 8.05 | 0.02 |

| *Enzyme | No. of acidic residues | No. of basic residues |
|---|---|---|
| VDE | 20 | 15 |
| CVDE | 79 | 84 |
